# Post-synaptic facilitation and network dynamics underlying stimulus-specific combination sensitivity

**DOI:** 10.1101/2025.08.25.672194

**Authors:** Zeina Merabi, Arij Daou

**Affiliations:** Neurophysiology and Computational Neuroscience Group, Biomedical Engineering Program, American University of Beirut, Lebanon

**Keywords:** combination sensitivity, temporal combination-sensitive neurons, temporal integration, spectral selectivity, complex auditory processing, biophysical modeling, neural circuits, postsynaptic facilitation, postsynaptic priming, coincidence detection

## Abstract

Combination-sensitive neurons (CSNs) integrate multiple stimulus features to generate behaviorally meaningful responses. While such neurons are well studied in fast timescale systems such as bat echolocation, the mechanisms enabling extended temporal integration in species like songbirds remain poorly understood. Using a computational model, we show that syllable-specific neurons in the songbird auditory system function as coincidence detectors whose selectivity depends on both post-inhibitory facilitation and persistent network activity. This persistence serves as a “memory trace”, allowing precise association of sequential syllables across hundreds of milliseconds. We propose that this dynamic interplay between intrinsic neuronal properties and convergent synaptic inputs gives rise to a higher-order “meta-combination sensitivity” enabling the auditory system to transform discrete acoustic events into temporally extended percepts. Our findings provide a mechanistic framework that bridges theories of coincidence detection with longer-timescale working memory, highlighting the importance of distributed network mechanisms for auditory temporal coding.

## INTRODUCTION

Sensory systems do not simply transmit information from the periphery to the brain; rather, they actively transform incoming signals to create meaningful representations of the external world. This transformation is essential for guiding behavior, allowing organisms to detect, interpret, and respond to complex environmental stimuli. A key aspect of this process is the extraction of behaviorally relevant information from peripheral inputs (Gomez-Ramirez, Hysaj et al. 2016). As sensory information ascends through processing hierarchies, neurons often exhibit selectivity for specific combinations of stimulus features (Carlson and Kawasaki 2006, Wenstrup, Nataraj et al. 2012). These so-called combination-sensitive neurons (CSNs) provide unique information that cannot be captured by single stimulus features alone (Margoliash and Fortune 1992, Doupe 1997, Medvedev, Chiao et al. 2002, Carlson and Kawasaki 2008). For example, the visual system integrates attributes like shape, size, color, and motion, and certain cortical neurons display striking selectivity for complex stimuli such as faces or objects (Fujita, Tanaka et al. 1992, Logothetis and Sheinberg 1996, Priebe and Ferster 2012).

Auditory systems also exhibit CSNs across species, including frogs (Mudry, Constantine-Paton et al. 1977, Fuzessery and Feng 1983), songbirds (Margoliash 1983, Takahashi and Konishi 1986, Margoliash and Fortune 1992, Lewicki and Konishi 1995, Lewicki 1996, Doupe 1997, Theunissen and Doupe 1998, Mooney, Rosen et al. 2002), bats (Suga, O’Neill et al. 1978, Suga, O’Neill et al. 1983), mice (Hofstetter and Ehret 1992), cats (Sutter and Schreiner 1991) and primates (Kanwal and Rauschecker 2007). In bats, delay-tuned neurons integrate pulse-echo pairs over millisecond timescales, serving as coincidence detectors crucial for echolocation (Wenstrup, Nataraj et al. 2012). Unlike bats, songbirds exhibit syllable-specific combination-sensitive neurons in HVC that integrate acoustic information over hundreds of milliseconds, enabling complex vocal communication and song discrimination (Margoliash and Fortune 1992, Lewicki and Konishi 1995, Mooney 2000, Mooney, Rosen et al. 2002). The bat’s combination sensitive behavior involve integrating information over ∼4 ms for delay-tuned neurons (Wenstrup, Nataraj et al. 2012, Suga 2015) and ∼20 ms for duration-tuned neurons (Aubie, Sayegh et al. 2012), while in songbirds this behavior is extended along a range of several hundreds of milliseconds (∼ 235 ± 73 ms) for a pair of syllables (Margoliash and Fortune 1992).

The extended temporal integration required in songbirds raises important questions: how do neurons preserve information across syllables, associate inputs with high temporal precision, and generate appropriate behavioral responses? In particular, how do syllable-specific CSNs manage to maintain the carried information from the first syllable ‘intact’ (Drew and Abbott 2003), prepare for the release and active association with the second syllable in an extremely precise timely manner over extended durations (Margoliash and Fortune 1992) and successfully generate an appropriately guiding behavior? One view is that such processing arises not solely from single-cell mechanisms but from dynamic, network-level transformations of neural activity over time (Karmarkar and Buonomano 2007, Paton and Buonomano 2018).

Building on this perspective and to address these challenges, we developed a time-dependent computational model to test how intrinsic neuronal properties and network dynamics contribute to extended temporal integration in combination-sensitive circuits. While our current investigations focus on songbirds, we contend that combination sensitivity is a ubiquitous neural mechanism with broad implications, applicable across diverse species and sensory systems where it plays a crucial role. Our results show how intrinsic neuronal properties, synaptic dynamics and upstream circuit mechanisms can endow CSNs with the capacity for precise temporal integration. We demonstrate that syllable-specific CSNs act as critical relay sites, integrating convergent inputs to function as high-order coincidence detectors. Within a narrow temporal window, they combine precisely timed network signals, a process shaped by post-inhibitory facilitation and sustained upstream activity that effectively maintains information between syllables. Our network-centric view provides a mechanistic account for how auditory circuits bridge discrete sensory events to construct coherent, temporally extended percepts, consistent with experimental observations across species.

## RESULTS

To evaluate how syllable-specific CSNs achieve selective integration over extended delays, we began by examining the canonical mechanisms proposed for two-tone combination processing in the auditory system. These include: (i) **temporal summation**, where successive subthreshold excitatory inputs combine to drive the neuron above firing threshold (Fricker and Miles 2000, Prescott and Sejnowski 2008), (ii) dual **post-inhibitory rebound (PIR)**, where two inhibitory inputs interact with the intrinsic properties of the CSN, particularly with the low-threshold T-type calcium channels, to produce a rebound burst in the CSN (Wenstrup, Nataraj et al. 2012, Kopp-Scheinpflug, Sinclair et al. 2018, Sun, Zhang et al. 2020); and (iii) **postsynaptic facilitation**, the most widely recognized mechanism, in which an initial inhibitory input primes the CSN by de-inactivating calcium channels and establishing a transient window of heightened excitability that enables a subsequent excitatory input to trigger robust spiking (Margoliash 1983, Lewicki and Konishi 1995, Mooney 2000, Mooney, Rosen et al. 2002, Dodla, Svirskis et al. 2006, Wenstrup, Nataraj et al. 2012, Dodla 2015).

While each of these mechanisms can account for short-timescale combination sensitivity, it remains unclear whether they are sufficient to explain the extended temporal integration observed in songbirds. We therefore developed computational models to test these scenarios systematically, and then extended them into a biologically realistic network framework that incorporates upstream dynamics. This approach enabled us to identify the limits of simplified motifs and to uncover how network-level mechanisms generate the temporally precise, syllable-pair sensitivity characteristic of songbird auditory circuits.

### Simple Network-Level Combinations

We started by developing a testable network model for each of these three mechanisms to evaluate the extent to which they would explain the observed neuronal behavior in songbirds, particularly for syllable-pair combination sensitivity, represented as syllables A and B. Figure 1A-C illustrate the network structure for each connectivity pattern described above. In particular, the network structure where the CSN receives two excitatory (Figure 1A) or two inhibitory (Figure 1B) inputs, tuned to syllables A and B and the case where the CSN receives an inhibitory input from syllable A and an excitatory input from syllable B (Figure 1C).

**Figure 1.**
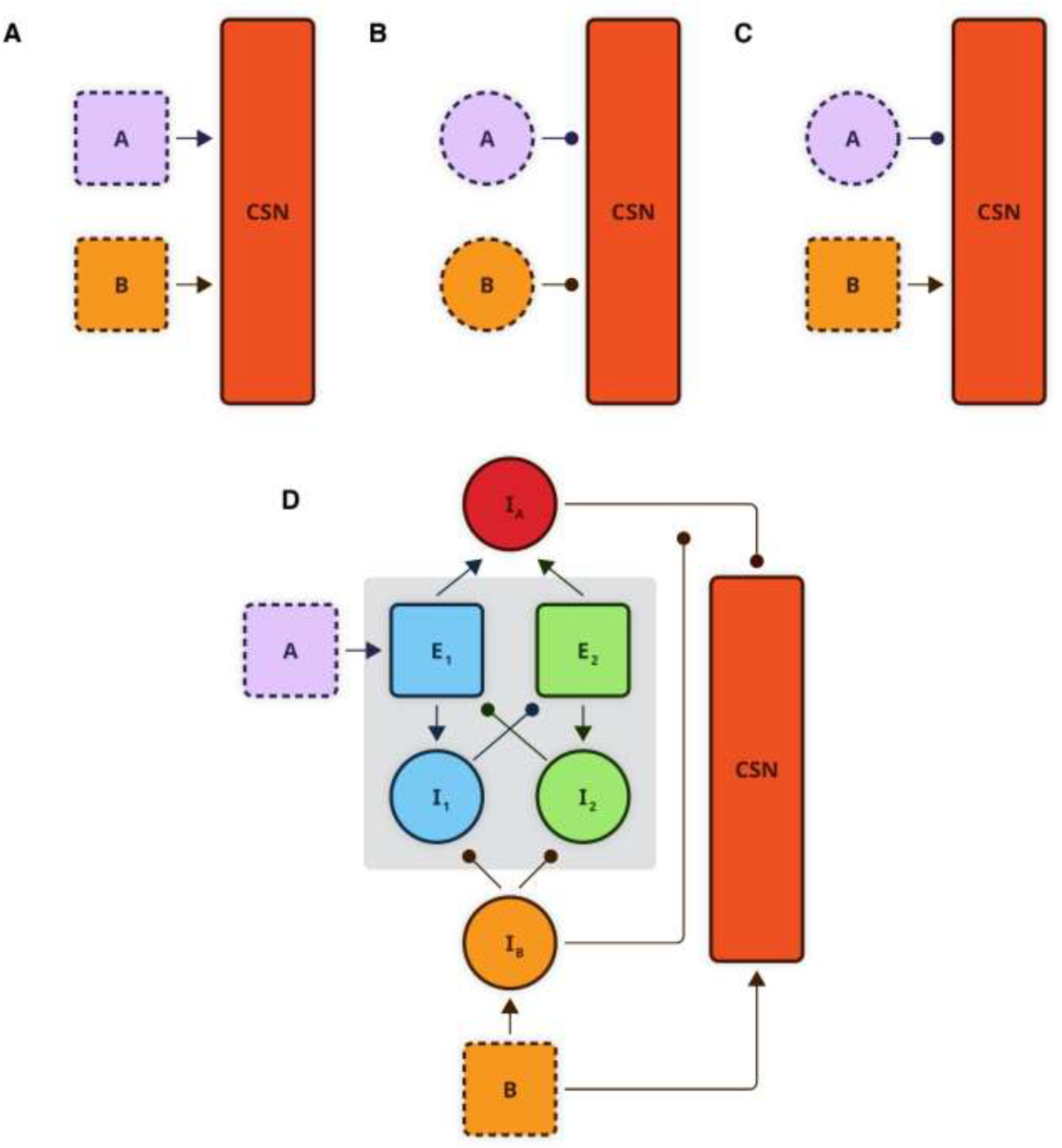
Network connectivity structures representing different mechanisms for a two-stimulus synaptic integration, with panels **(A-C)** illustrate simple network scenarios extended from existing literature, while panel **(D)** illustrates our novel biologically realistic network model. **(A)** Temporal Summation: Convergent excitatory inputs from neurons A (pale lavender) and B (yellow) onto the combination-sensitive neuron (CSN, orange). **(B)** Dual Post-Inhibitory Rebound (PIR): Convergent inhibitory inputs from interneurons A and B onto the CSN. **(C)** Postsynaptic Facilitation: Convergence of an inhibitory (interneuron A) and excitatory (neuron B) input onto the CSN. **(D)** Postinhibitory facilitation underlying combination sensitivity and incorporating temporal delay. Gray shaded region depicts the temporal delay loop incorporating two excitatory (E1 and E2) and two inhibitory (I1 and I2) neurons. The network also includes interneurons I_A_ (red circle) and I_B_ (yellow circle). Solid arrowheads indicate excitatory synaptic transmission; rounded arrowheads indicate inhibitory synaptic transmission. Excitatory neurons are represented by squares and excitatory inputs by solid arrowheads, while inhibitory neurons are represented by circles and their inputs by rounded arrowheads. The CSN is represented by a rectangle.

For each of the simple network scenarios shown in Figure 1, we simulated the underlying network dynamics under different conditions. For all simulations throughout this paper, we refer to syllable A as the first stimulus (or S1) and syllable B as the second stimulus (or S2), for greater generality. Starting with the temporal summation scenario (Figure 1A), the firing patterns of the neurons representing S1, S2 and CSN are shown in Figure S2, when both stimuli arrive sequentially (Figure S2-A), when the firing of S2 is delayed by 50 ms from the offset of S1 (Figure S2-B), when the temporal order of arriving stimuli is reversed (Figure S2-C), and when the CSN receives two identical excitatory stimuli (for example, representing identical spectral features when syllable A is repeated, Figure S2-D). When both stimuli arrive in succession, the first input elevates the CSN’s membrane potential to a subthreshold level, increasing its excitability and enabling the generation of a facilitative response triggered by the second input that arrives within a strict temporal window (Figure S2-A). A short delay before the arrival of the second stimulus results in a combination failure for this scenario (Figure S2-B), because even a delay as short as 50 ms is sufficient to bring neuron A’s membrane potential to resting state, rendering S2’s effect useless to trigger a suprathreshold response in the CSN. More importantly, this network seems to be insensitive to the temporal order of syllables, in that reversing the order of stimuli yields successful behavior (Figure S2-C). Spectral sensitivity was also absent since the CSN is stimulated by the summation of subthreshold inputs regardless of their spectral selectivity (Figure S2-D). We performed similar simulations for the second scenario (Figure 1B), where the CSN receives dual inhibitory inputs (Figure S3). The resulting network behavior closely resembles that observed in the first scenario. These findings render such simplified network structures incapable of explaining the whole mechanism of syllable combination sensitivity. It is noteworthy that we are not rejecting these neuronal mechanisms outright, as they are effective in the absence of delay. Instead, we are showing that the extended temporal delays and strict temporal/spectral sensitivity characteristic of syllable-specific combination-sensitive neurons are not accounted for by such a simplified three-cell network structure.

In this study, we focus on the postsynaptic facilitation mechanism (Figure 1C), in which the CSN receives spectrally distinct inhibitory and excitatory inputs. This mechanism, referred to as postsynaptic facilitation or priming, has been widely proposed to underlie syllable combination in songbirds (Margoliash and Fortune 1992, Lewicki and Konishi 1995, Mooney, Rosen et al. 2002). To assess its dynamics, we examined the simple network’s response under four conditions: when the excitatory input coincides with the offset of S1 (Figure 2A), when S2 arrives too early before S1 ends (Figure 2B), when the arrival of S2 is delayed by 50 ms (Figure 2C), and when the temporal order of syllables is reversed (Figure 2D).

**Figure 2.**
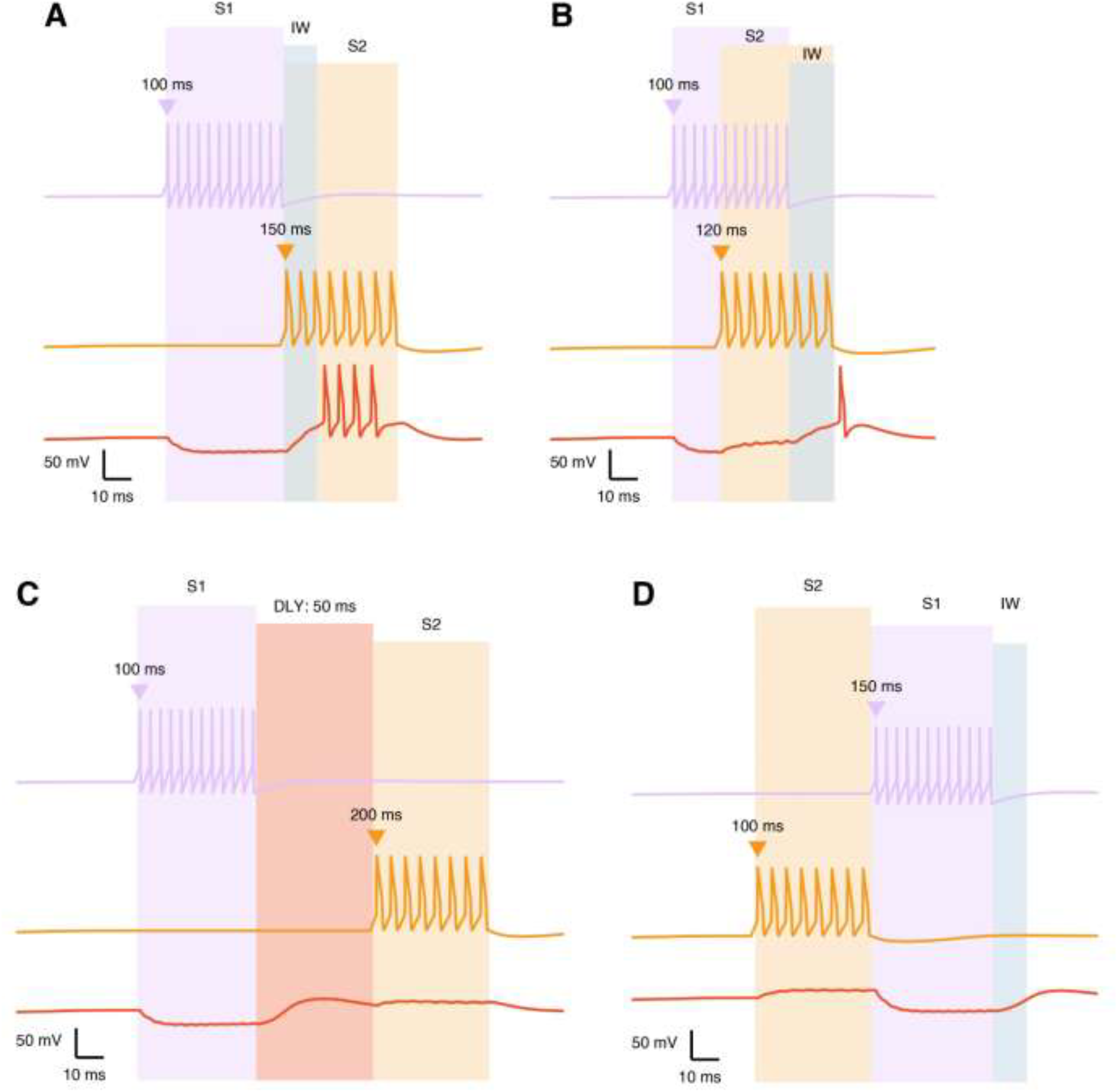
Simple network dynamics for the postsynaptic facilitation scenario. Firing patterns for an inhibitory interneuron responding to the first stimulus (S1, pale lavender trace), an excitatory neuron responding to the second stimulus (S2, yellow trace) and the CSN (orange trace) are shown under different conditions **(A-D).** The onsets of S1 (pale lavender triangle) and S2 (yellow triangle) are varied across the different panels, as well as their respective durations depicted by the shaded regions of the same colors. **(A)** S1 (50 ms) followed by S2 (50 ms) immediately results in successful integration at the CSN level generating a post inhibitory rebound burst. An integration window (IW, light blue shaded area) refers to a temporal window of CSN’s heightened excitability. **(B)** S2 arriving before the end of S1 leads to integration failure. **(C)** Temporal delays (DLY: shaded soft red area, 50 ms) outlasting the CSN’s integration window fails to associate the CSN. **(D)** Reversed stimulus order fails to associate the inputs, showing a dissociated response.

In the case of successful integration (Figure 2A), interneuron A fires at the onset of S1 (t = 100 ms), delivering an inhibitory input to the CSN that persists for the duration of the stimulus (50 ms, shaded pale lavender area). This hyperpolarization primes the CSN, which begins a rebound response at stimulus offset (t ∼ 150 ms) due to its T-type Ca^2+^ and H-currents (see Methods). The resulting rebound window, lasting for a few milliseconds (∼16 ms), defines the CSN’s integration window (IW, shaded light blue), during which its intrinsic excitability is enhanced. The larger the T-type Ca^2+^ conductance, the shorter the integration window. When S2 activity overlaps with this window, the excitatory input coincides with the rebound depolarization, driving the CSN above threshold and producing a maximal facilitatory response that signifies successful syllable combination.

Temporal variation in the arrival of the second stimulus disrupts effective integration at the CSN. When S2 arrives prematurely, before the offset of S1 (e.g., t = 120 ms, Figure 2B), or arrives outside the integration window, for instance, with a 50 ms delay (Figure 2C), the CSN fails to generate a combined response and instead exhibits two temporally dissociated subthreshold events. Similarly, reversing the temporal order of the stimuli abolishes coincidence detection, as the CSN does not reach spiking threshold under this condition (Figure 2D). One of the main limitations of simplified network models, is that they primarily focus on postsynaptic events taking place at the CSN level, which cannot fully explain behaviors that operate on larger timescales. We suggest that delays are dynamically managed by an upstream network level interaction, presynaptic to CSN.

### Time Encoding Network

#### Temporal Delay and Integrations

To address the complexities of combination-sensitive behavior in songbirds, we next examined how temporal information is encoded at the network level and how such encoding supports the association of syllables across extended delays. To this end, we expanded the simplified network model by incorporating a temporal delay loop capable of generating all the biologically realistic mechanisms of combination sensitivity (Fig. 1D). This loop sustains reverberatory activity that allows the network to encode a “memory trace” of the first stimulus. It consists of two excitatory neurons (E_1_ and E_2_, blue and green squares), and two inhibitory interneurons (I_1_ and I_2_, blue and green circles). Blue colored neurons (E_1_ and I_1_) initiate the loop’s activity upon the onset of the first stimulus, while green colored neurons (E_2_ and I_2_) propagate this activity, sustaining a “memory trace” of the stimulus after its offset. A feedback input from I_2_ to I_1_ encloses the loop, allowing this activity to span an inter-stimulus duration. Excitatory input from E_1_ and E_2_ also drive an intermediate interneuron (I_A_, red circle), which lies outside the loop and relays important inhibitory signals to the CSN (orange rectangle), as will be described next. In parallel, a second interneuron (I_B_, yellow circle), activated by the B-selective excitatory neuron, provides inhibitory input to I_1_ and I_2_, thereby regulating loop activity. Eventually, the CSN integrates direct excitatory input from the B-selective neuron with inhibitory input from I_A_, establishing the basis for temporally precise combination sensitivity.

The activity patterns that this extended network exhibits are shown in Figure 3A, which shows the spike trains of all neurons in response to a 150 ms delay (shaded soft red area) between the offset of the first stimulus and the onset of the second. When S1 is presented (pale lavender triangle), the A-selective excitatory neuron fires for 50 ms (pale lavender shaded area). Although it doesn’t connect directly to the CSN, it initiates the integration process by activating E_1_, thereby engaging the temporal delay loop. E_1_ propagates its activity along two pathways: The first pathway is an exterior one, where it excites the inhibitory interneuron I_A_. I_A_ in turn sends an inhibitory input to the CSN, effectively hyperpolarizing its membrane potential. The second pathway remains within the loop, where E_1_ excites interneuron I_1_, which in turn suppresses E_2_. When I_1_ activity ceases, E_2_ is able to escape its inhibition and generates a rebound burst mediated by its T-type Ca^2+^ and H-currents. The rebound that E_2_ generates maintains loop activity and provides a second inhibitory drive to the CSN via I_A_ (dashed line with solid arrowheads from E_2_ through I_A_ to the CSN), thereby preventing premature post-inhibitory rebound at the CSN (before the second stimulus arrive). At the same time, it plays a coding function, where I_2_ reactivates E_1_ through feedback inhibition, sustaining a reverberatory cycle that preserves a memory trace of S1 throughout the inter-stimulus interval. As shown in Figure 3B, this mechanism is robust across a range of delays, with I_A_ firing continuously during the delay period (due to the excitatory inputs from E_1_ and E_2_), ultimately maintaining CSN hyperpolarization.

**Figure 3.**
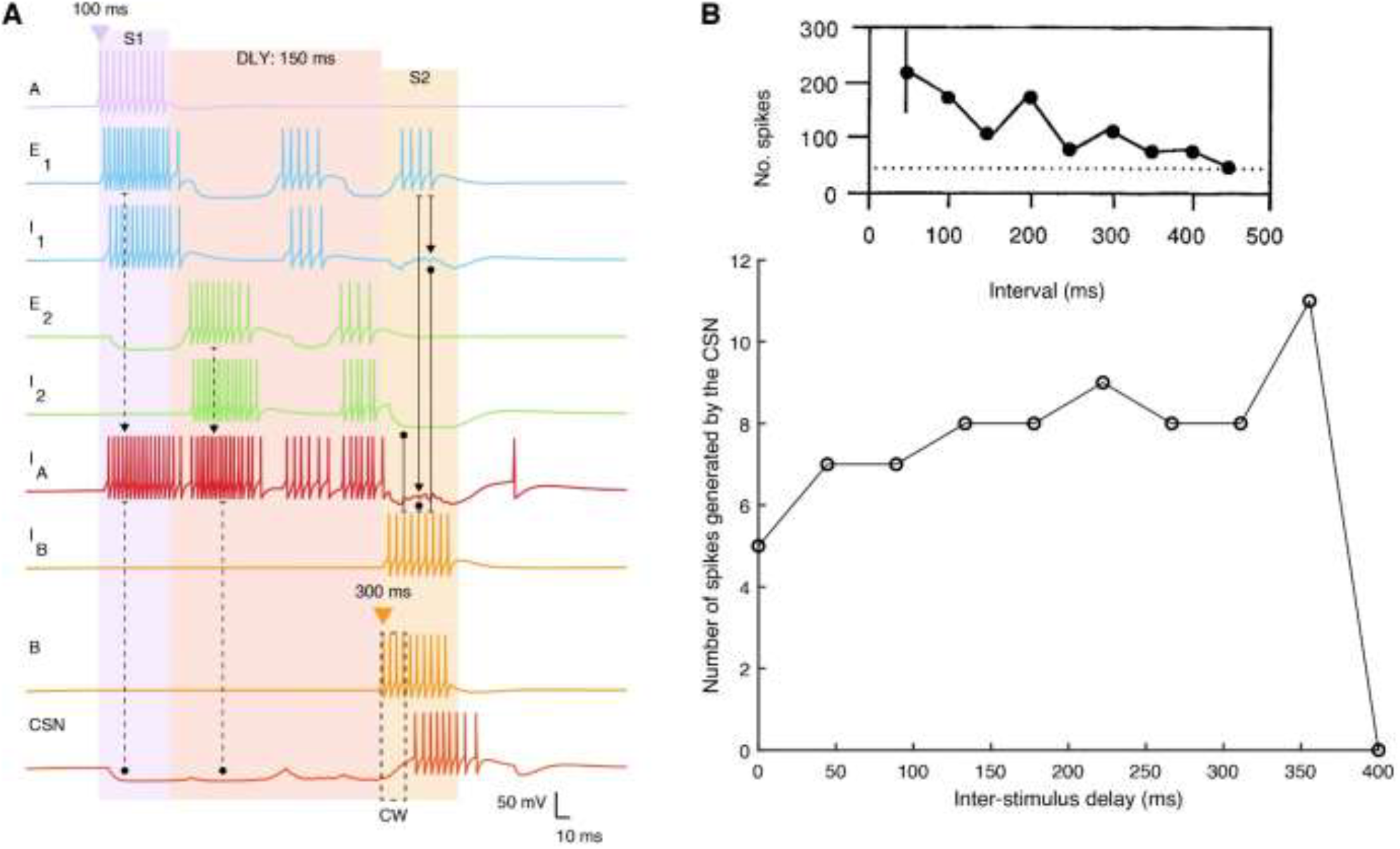
Combination-sensitive behavior maintained along extended delay durations. **(A)** Firing patterns of neurons in the suggested network depicting the dynamics in response to S1 and S2, separated by a 150 ms delay (DLY: soft red area). Voltage traces are color-coded according to their representation in the schematic of Figure 1D, with neuron names indicated on the left. Dashed and solid arrows between different traces highlight key events along the temporal propagation of activity. Coincidence window (CW) at the CSN trace is highlighted by a dashed rectangle. Excitatory synaptic connections are represented by solid arrowheads, and inhibitory synaptic connections by rounded arrowheads. **(B)** Number of spikes generated by the CSN as a function of inter-stimulus delay. Inset displays experimental data from (Margoliash and Fortune 1992), showing a similar trend with our obtained results.

The loop is terminated at the onset of S2 (yellow triangle), which signals the end of the delay period and activates a B-selective excitatory neuron. This neuron plays two simultaneous functions: it directly excites the CSN and triggers dis-inhibitory actions through the recruitment of interneuron B (I_B_), which delivers inhibitory inputs to I_1_, I_2_ and I_A_. These inhibitory inputs counteract concurrent excitatory drives within the loop, suppressing their activity and thereby terminating the inhibitory influence of I_A_ on the CSN. Once the inhibition is lifted, the CSN undergoes a post-inhibitory rebound, creating a brief temporal window, referred to as a coincidence window (CW, dashed rectangle) of heightened excitability (∼27 ms in this simulation). The duration of this window is dependent on a balanced strength of the CSN’s intrinsic characteristics, allowing it to function as a critical network-level transformation point (as will be illustrated next). Notably, this window is not dynamically extensible, and have a limited temporal capacity. During this window, the coincident excitatory input from the B-selective neuron aligns with the rebound depolarization, driving the CSN to fire maximally and producing successful stimulus integration.

#### Dual integration windows: A dynamic shift

Our network simulations reveal a dynamic shift in temporal coding strategies, demonstrating two distinct, yet interdependent, integration windows crucial for spectrotemporal integration and temporal encoding. The first is an extended “integration period” which can span several hundreds of milliseconds (Margoliash 1983), reflecting the inter-stimulus delay duration. This window is mediated by reverberatory loop activity at the I_A_ interneuron level, and is mediated at the CSN level by asynchronous excitatory inputs from E_1_ and E_2_. The second is a short “coincidence window” lasting only tens of milliseconds, during which the CSN integrates excitatory and rebound inputs to generate a combined response.

To quantify the temporal limits of these mechanisms, we varied the inter-stimulus delay duration from 0 to 400 milliseconds in ∼20 milliseconds steps and measured CSN spiking (Figure 3B). As delays increased, spike counts fluctuated before declining sharply, with activity abolished at ∼400 ms. This pattern broadly aligns with the experimental data from Margoliash and Fortune (1992). However, whereas their recordings showed a gradual monotonic decline in spiking with increasing delay, our model exhibited an initial increase up to ∼350 ms followed by a rapid drop. This discrepancy likely reflects differences in experimental context, as Margoliash investigated neurons responding to syllable sequences (syllables 2” - 6”), whereas our model focuses on pairwise syllable integration governed by precisely orchestrated interactions between upstream network dynamics and CSN intrinsic properties.

The sharp abolishment of responses near 400 ms is consistent with the paradoxical gating of T-type calcium channels (Leresche, Hering et al. 2004). While hyperpolarization typically promotes deinactivation of T-channels (Huguenard 1996, Perez-Reyes 2003, Sun, Zhang et al. 2020), prolonged hyperpolarization can trigger phosphorylation-mediated inhibition, paradoxically inhibiting the T-current and reducing its availability (Leresche, Hering et al. 2004). Therefore, an optimal T-current isn’t generated after prolonged hyperpolarization, but rather after transient ones (Dodla, Svirskis et al. 2006), where the balance between dephosphorylation and deinactivation kinetics is favorable. Thus, extended inhibitory periods (>= 350 - 400 ms) may paradoxically suppress rebound excitability in the CSN despite continued inhibition. Although direct evidence for this mechanism in zebra finches is limited, kinase-mediated regulation of ion channels has been implicated in the song system plasticity (Aronowitz, Kirn et al. 2022).

Another possible explanation could be imposed by the dynamics of the network’s delay loop itself. At extended delays, feedback inhibition-driven rebounds in excitatory neurons (E_1_, E_2_) attenuate (Figure 4A), potentially due to T-channel phosphorylation effects. This reduces the excitatory drive to I_A_, weakening its inhibitory control over the CSN. Consequently, fewer T-type channels are deinactivated in the CSN, weakening its post-inhibitory rebound and limiting spiking at longer delays.

**Figure 4.**
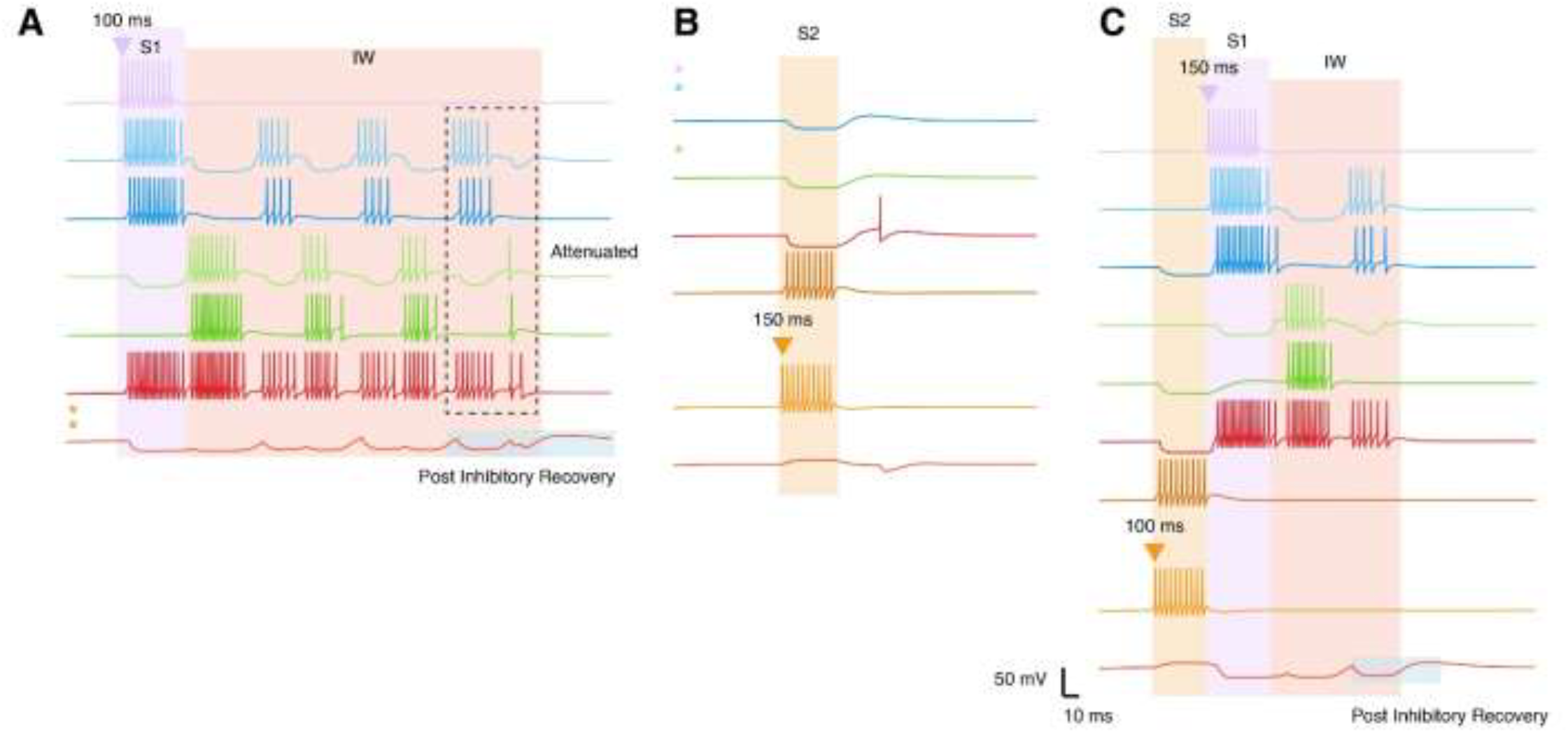
Network simulations of unsuccessful combination behavior. **(A)** Response to S1 presentation solely. The onset of S1 (pale lavender triangle) initiates a cascade of firing (pale lavender shaded area). S1 offset, characterized by a ping-pong firing, defines the integration window (IW, soft red area). As the IW progresses, the firing patterns within the delay loop attenuate (dashed rectangle), resulting in a gradual decrease of the inhibitory drive leading to post-inhibitory recovery (overlapping light blue shade around the CSN trace). **(B)** Response to S2 presentation solely (yellow triangle represents onset of S2). **(C)** Response to reversed stimulus order (S2 preceding S1, with a 0 ms delay). The CSN exhibits a gradual post-inhibitory recovery towards the end of the IW, rather than a sharp release leading to robust spiking. Untrigerred voltage traces are excluded, replaced with appropriate color-coded asterisks.

To validate these dynamics, we further examined extreme cases of at 0 and 300 ms delays (Figure S4). Notably, the duration of the coincidence window was relatively constant in all conditions. At 0 ms delay (Figure S4-A), both integration windows were activated immediately, producing robust coincidence detection. At 300 ms delay (Figure S4-B), the network preserved S1 activity through an elongated integration window, successfully transitioning to coincidence detection at the onset of S2. Together, these findings highlight the cooperative operation of two temporally distinct windows that enable CSNs to encode syllable associations across both short and extended timescales.

#### Unsuccessful Integration

Combination-sensitive neurons are known to exhibit an all-or-none response, whereby combination-sensitive activity is successfully generated only when both syllables are presented in the correct order. The above findings highlight the important roles of both stimuli onsets and offsets, particularly how their sequential order governs the dynamic transition from the extended temporal integration window to the brief coincidence detection window. This sequential ordering is key for successful stimulus combination. Here, we further elaborate on this principle by examining how the isolated presentation of a single stimulus (syllable), or their reversed order, fails to elicit the appropriate behavioral output (Figure 4).

When only the first stimulus (S1) is presented (Figure 4A), its onset and offset trigger upstream network activity that opens an extended integration window (∼370 ms, shaded soft red area). During this period, the sustained activity of I_A_ maintains inhibition of the CSN, but this inhibitory drive gradually weakens as rebound activity in E_1_ and E_2_ attenuates across repeated cycles of hyperpolarization (dashed rectangle). Reduced I_A_ firing reduces CSN hyperpolarization, allowing gradual recovery without producing a robust rebound burst (shaded light blue area). Thus, while S1 offset initiates temporal integration, it is insufficient on its own to drive CSN spiking, highlighting the role of the CSN as a coincidence detector (Margoliash 1983, Lewicki and Konishi 1995, Doupe 1997, Prather and Mooney 2008, Wenstrup, Nataraj et al. 2012, Suga 2015, Medina and Margoliash 2024).

Similarly, S2 presented in isolation fails to evoke CSN firing (Figure 4B). Its excitatory input alone produces only subthreshold depolarization in the CSN, and without upstream activity the disinhibitory role of I_B_ is effectively lost. Reversing the temporal order of syllables (S2 preceding S1) also abolishes combination, yielding dissociated responses reminiscent of the simplified network model (Figure 4C). Together, these emphasize the unidirectional transition from integration (initiated by the offset of S1) to coincidence detection (triggered by the onset of S2) as a fundamental requirement for effective association of stimuli.

To ensure that network responses were not due to simple facilitation, we tested repeated presentation of the same stimulus. Both S1-S1 (Figure S5-A) and S2-S2 (Figure S5-B) yielded responses equivalent to isolated presentations, confirming spectral selectivity and ruling out nonspecific summation effects. These findings emphasize that combination sensitivity is an emergent network property, shaped by the precise coordination of excitation, inhibition, and temporal windows, rather than by single-cell mechanisms alone.

We next examined the role of intrinsic membrane conductances in shaping CSN responses. Under normal conditions (default scenario, same parameters for CSN’s conductances as in Figure 3), the CSN neuron elicited a six-spike burst (Figure 5, red trace). Eliminating the H-current (*g*_*h*_= 0, green trace) produced stronger hyperpolarization during integration, but the CSN still generated a rebound burst due to 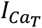. In contrast, eliminating the T-type calcium current (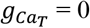, blue trace) while keeping *g*_*h*_intact abolished rebound bursting, leaving only a small subthreshold depolarization due to *I*_*h*_, highlighting the important role of 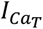in modulating the CSN’s bursting activity. Reversing syllable order disrupted this synaptic-intrinsic interplay, preventing successful integration (Figure S6), consistent with postsynaptic priming models (Mooney, Rosen et al. 2002).

**Figure 5.**
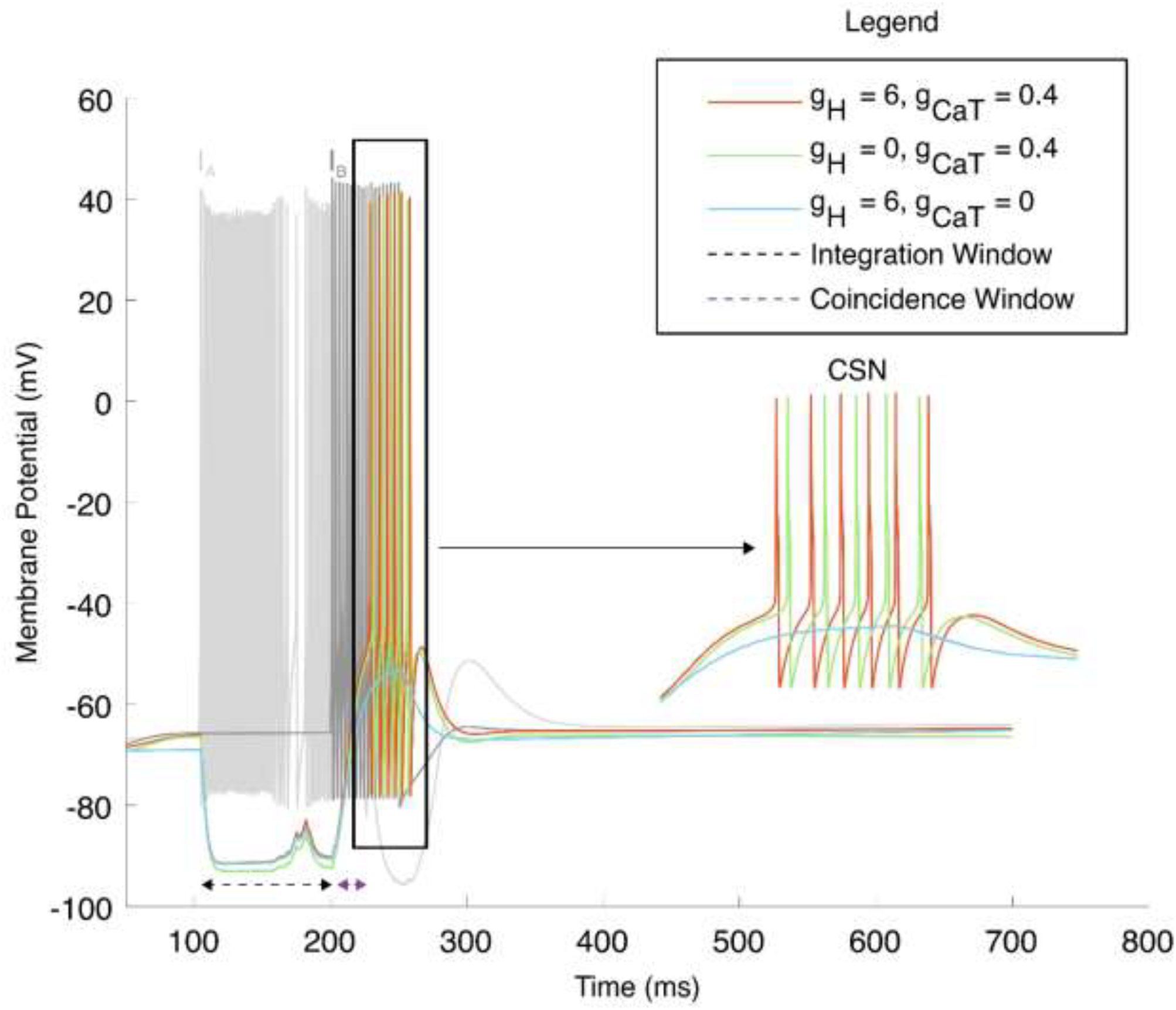
CSN post-synaptic priming dynamics. The figure illustrates the membrane potential traces of interneuron I_A_ (light grey), neuron B representing S2 (darker grey), and the combination-sensitive neuron (CSN, colored traces) during a simulation where Stimulus 1 is followed by Stimulus 2 with 50 ms delay. The interneuron I_A_’s firing defines an integration window (black dashed line, a 100 ms integration in this case), followed by a brief coincidence window (purple dashed line, ∼27 ms) triggered by neuron B’s activity. The red trace (default conductances, as in Figure 3) shows successful integration with 6 spikes rebound burst. The green trace (blocking I_h_while keeping 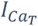intact) exhibits a slightly delayed and reduced 5-spike burst, with more hyperpolarization exhibited during the integration window due to the lack of I_h_. The blue trace (blocking 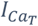 while keeping I_h_intact) matches the red trace during the integration window, exhibiting a slight rebound depolarization due to I_h_, but fails to reach the threshold level to produce any postinhibitory burst activity due to the lack of 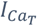.Inset shows a zoomed version of the rebound region (enclosed by the black rectangle).

To further explore the cooperative roles of 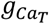 and *g*_*h*_in gating the integration process, we systematically varied their conductances (Figure 6A). The resulting heatmap revealed a step-like pattern of spike counts, reflecting their complementary interaction. For example, low 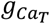 (0.4 nS) combined with minimal *g*_*h*_(0.1 nS) evoked only 6 spikes (highlighted white square). Increasing either conductance enhanced CSN spiking, with higher *g*_*h*_extending burst duration and higher 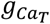 amplifying rebound initiation. This demonstrates that 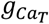 primarily drives burst onset/initiation, while *g*_*h*_sustains depolarization, together shaping the CSN’s ability to generate prolonged firing.

**Figure 6.**
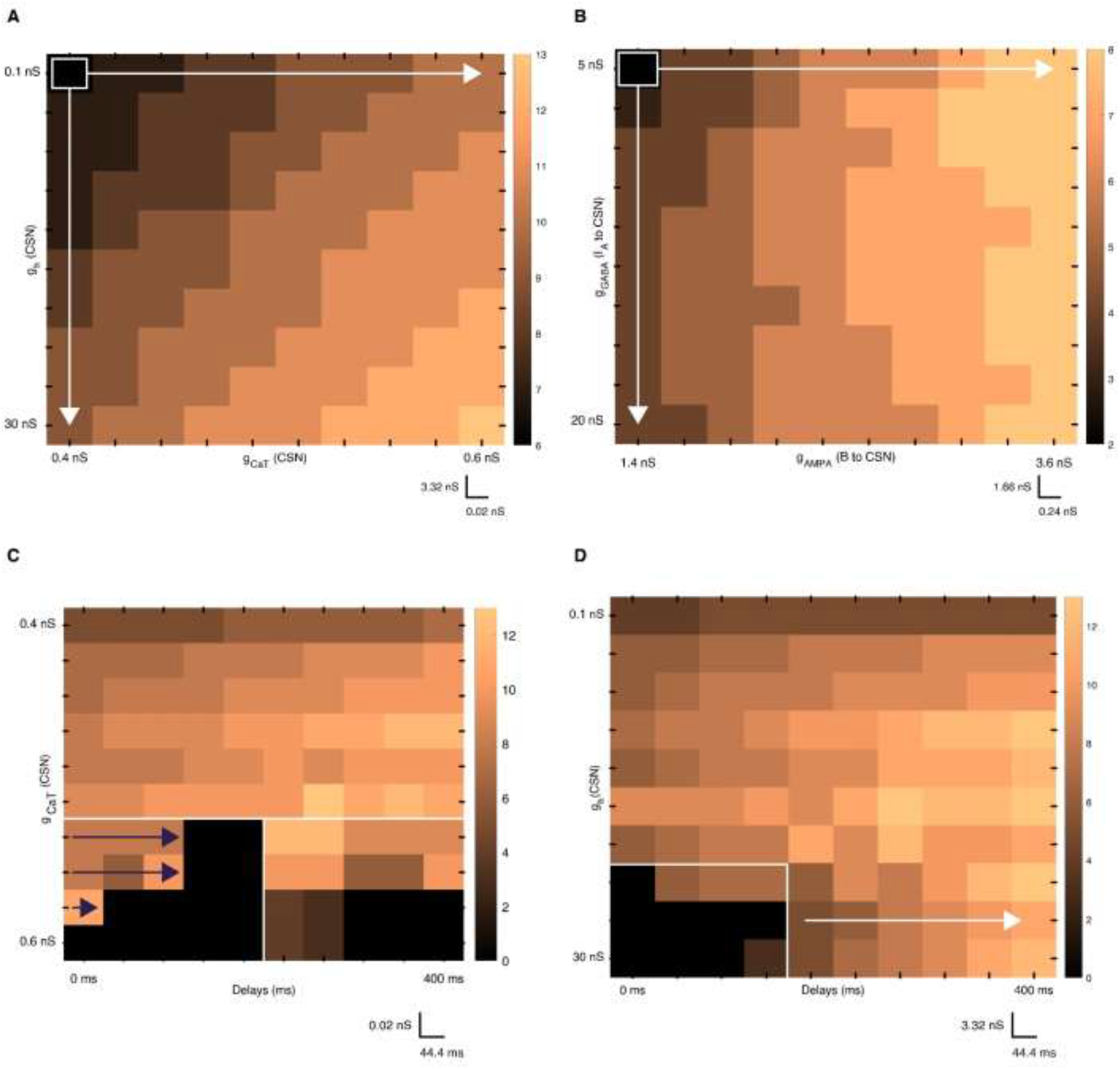
**(A-B)** Intrinsic and synaptic modulations of the CSN firing. **(A)** Intrinsic determinants of postsynaptic priming in the CSN: Heatmap illustrates the relationship between 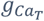 (x-axis: 0.4-0.6 nS in 0.02 nS increments) and g_h_(y-axis: 0.1-30 nS in 3.32 nS increments) and its impact on the CSN spiking behavior (spike count) while keeping all other parameters intact as in Figure 3. **(B)** Synaptic determinants of postsynaptic facilitation in the CSN: Heatmap illustrates the relationship between the excitatory g_AMPA_ (B to CSN) (x-axis: 1.4-3.6 nS in 0.24 nS increments) and the g_GABA_ (IA to CSN) (y-axis: 5-20 nS in 1.66 nS increments) and its impact on the CSN spiking behavior, while keeping all other parameters fixed as in Figure 3. **(C-D)** Interplay of temporal delay and intrinsic neuron properties in CSN integration. **(C)** Delays vs. CSN’s Intrinsic gCaT Conductance: This 10×10 heatmap shows CSN spike count as a function of inter-stimulus delay (x-axis: 0-400 ms) and gCaT conductance (y-axis: 0.4-0.6 nS). **(D)** Delays vs. CSN’s Intrinsic gh Conductance: This 10×10 heatmap displays CSN spike count as a function of inter-stimulus delay (x-axis: 0-400 ms) and gh conductance (y-axis: 0.1-30 nS). The color intensity of each pixel represents the number of spikes generated by the CSN for that specific parameter combination, with the corresponding spike count indicated by the color bar.

### Exploring the synaptics: Postsynaptic facilitation

We next examined how synaptic conductances shape CSN activity, focusing on the inhibitory input from I_A_ and the excitatory input from the B-selective neuron. Using an approach analogous to that applied for intrinsic currents, we systematically varied both synaptic conductances (*g*_*GABA*_ and *g*_*AMPA*_, see Methods) and quantified the resulting CSN spike output.

Figure 6B illustrates the interplay between the GABAergic and the glutamatergic inputs to the CSN, driven by the I_A_ and B-selective neurons, respectively. The heatmap shows a fairly regular horizontal gradient, with spike counts increasing as excitatory synaptic conductance rises, while variation in inhibitory synaptic conductance alone exerts a weaker effect. At the lowest values of both inputs (highlighted square), the CSN failed to exhibit successful combination. Increasing *g*_*GABA*_ while holding *g*_*AMPA*_ constant produced only modest spiking increases, reaching its maximum at the highest inhibitory conductance with minimal excitation (bottom of the first column). In contrast, progressively raising *g*_*AMPA*_ values led to a marked increase in spiking.

These findings raise two important key points. First, the limited influence of *g*_*GABA*_ alone suggests that inhibitory strength must be matched with sufficient intrinsic excitability, particularly from *g*_*CaT*_, to enable robust rebound firing (Figure 6A). Without this intrinsic support, stronger inhibition does not necessarily translate into greater facilitation. Second, the stronger effect of increasing *g*_*AMPA*_ highlights the decisive role of excitatory drive following inhibition, confirming its facilitatory contribution to postsynaptic integration. Together, these findings computationally validate postsynaptic priming as the central mechanism for combination sensitivity, highlighting the critical interplay between synaptic and intrinsic factors in shaping CSN responses.

#### The Interplay between the Two Integration Windows

Our simulations demonstrate the obligatory transition from temporal integration to coincidence detection renders the network highly sensitive to disruptions in either window. To understand the relationship between these two temporal windows, we examined key parameters characterizing each and analyzed their impact on the combination behavior, quantified by the number of CSN spikes. The temporal integration window was characterized by inter-stimulus delay, whereas the coincidence window was defined by intrinsic parameters, specifically 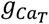 and *g*_*h*_. Two heatmaps (Figure 6C-D) illustrate the combined effects of these parameters on CSN firing

For the delay − 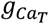 analysis (Figure 6C), the heatmap exhibited a horizontally stratified pattern with a cut-off near 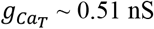 (white line). Above this threshold, CSN spiking activity generally increased with longer delays and higher conductance values. Along the delay x-axis, spike counts increased progressively, while along the conductance axis, results were more heterogeneous.

Mechanistically, these results suggest that sustained inhibition from I_A_ prolongs CSN hyperpolarization, with 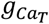 determining the neuron’s readiness to generate rebound depolarization following inhibition release. Higher 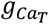 values enhance this rebound, supporting more reliable spiking.

Below the cut-off region, however, responses became irregular, with many parameter combinations yielding no spikes (dark-colored areas). Failures were particularly common at shorter delays (<180 ms, highlighted by the purple arrows), where increasing 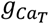 paradoxically reduced spike generation, indicating a delicate balance between the delay and conductance parameters to align the temporal integration and coincidence detection windows. At longer delays (>180 ms) with high 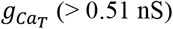 (> 0.51 nS), the parameter - response relationship became highly nonlinear and non-uniform, with alternating zones of strong and absent spiking, reflecting complex interactions between synaptic and intrinsic mechanisms.

The delay - *g*_*h*_analysis (Figure 6D) revealed similar trends. At short delays and high *g*_*h*_values, spiking was completely abolished (outlined region), because excessive depolarization prevented activation of the T-type Ca^2+^ current. For longer delays (traced by the white arrow), regions of maximal spiking emerged, suggesting that additional ion channel recruitment may sustain excitability even when *g*_*h*_is high.

Collectively, our results suggest that CSN responses are governed by a finely tuned interplay between intrinsic parameters, synaptic inputs, and temporal factors. The interaction of these factors defines the operating ranges of the integration and coincidence windows; reinforcing our hypothesis that the CSN actively shapes output patterns rather than serving as a passive relay of convergent inputs.

#### Stimuli Characteristics and the Generalizability of our Network

Finally, we run further analysis to assess the robustness of our network behavior in response to variations in stimuli characteristics, specifically stimulus duration and intensity. While preceding simulations used default values (50 ms duration, 200 pA intensity), we expanded the parameter space to include stimulus durations up to 200 ms and intensities up to 400 pA. This broader exploration allowed us to probe the network’s response to a wider range of syllable-like inputs, reflecting natural variability in temporal and spectral features that may be integrated by the CSN.

### Stimulus duration

We first examined stimulus duration using a single-parameter variation approach, systematically varying the duration of one stimulus while holding the other constant. With an interstimulus delay fixed at 50 ms, we explored durations from 10 to 200 ms for each input. As shown in Figure 7A, very brief presentations of the first stimulus (10 ms, pale lavender) were sufficient to initiate CSN firing, indicating that the network is sensitive to the occurrence of S1 rather than its precise duration. This is consistent with the network architecture, in which neuron A activates the temporal delay loop, effectively acting as a “wake-up” signal, without directly synapsing onto the CSN. Increasing S1 duration progressively raised spike output, indicating that, irrespective of the first stimulus’s duration, once the delay loop is activated, subsequent neuronal events unfold sequentially. In contrast, the CSN displayed greater sensitivity to the duration of the second stimulus. A minimum duration of 30 ms was required to elicit spiking, with spike counts increasing steeply to a plateau at durations >= 100 ms. This heightened sensitivity reflects the direct excitatory input from neuron B to the CSN, which amplifies output as S2 lengthens.

**Figure 7.**
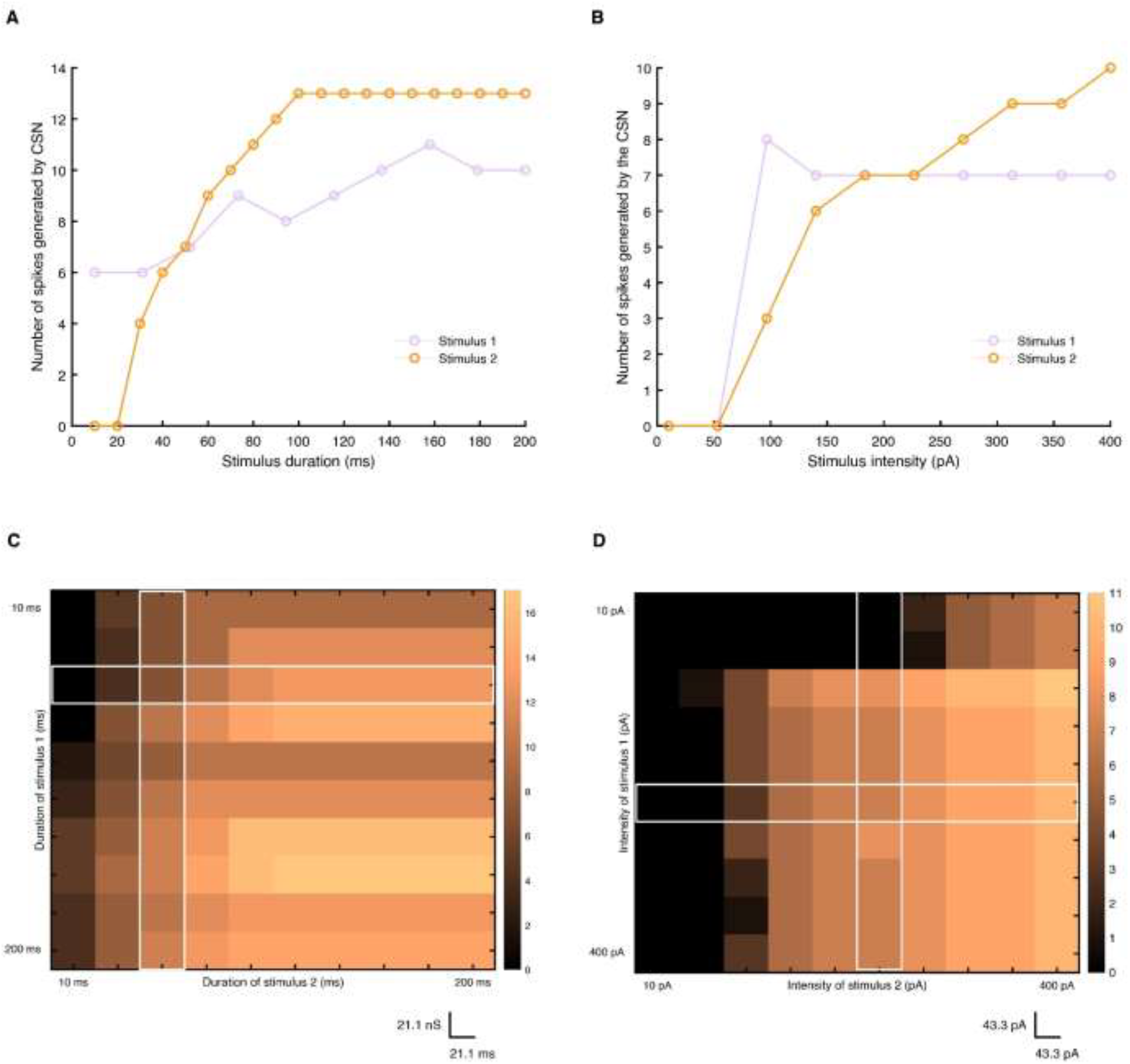
**(A-B)** Influence of stimulus characteristics on successful combination behavior (CSN spike count). **(A)** Line plots for S1 and S2 duration variation vs. CSN spike count. The purple line represents the effect of varying the duration of S1 and the yellow line represents the effect of varying the duration of S2. The x-axis represents stimulus duration (from 0 to 200 ms), and the y-axis represents the number of spikes generated by the CSN. **(B)** Line plots for S1 and S2 intensity variation vs. CSN spike count. The purple line represents the effect of varying the intensity of S1 and the yellow line represents the effect of varying the intensity of S2. The x-axis represents stimulus intensity (from 0 to 400 pA), and the y-axis represents the number of spikes generated by the CSN. **(C-D)** Generalizability of our network model and its response across varying stimulus characteristics (Duration and Intensity). **(C)** Stimulus duration variations: pixels of a heatmap (10×10) recording parameter combination values that illustrate the relationship between the duration of S1 and the duration of S2 on the firing response of the CSN; x-axis: 10-200 ms in 21.1 ms increments. **(D)** Stimulus intensity variations: pixels of a heatmap (10×10) recording parameter combination values that illustrate the relationship between the intensity of S1 and the intensity of S2 on the firing response of the CSN; x-axis: 10-400 pA in 43.3 pA increments. The color intensity of each pixel represents the number of spikes generated for each specific combination, with the corresponding spike count indicated by the color bar on the left.

To further generalize these results, we co-varied the durations of both stimuli across the full 10 to 200 ms range while maintaining default conductance parameters and a 50 ms interstimulus delay. The resulting heatmap (Figure 7C) confirmed that variations in S1 duration had relatively little effect on CSN activity (uniform intensity along the x-axis), whereas increasing S2 duration yielded a significant rise in spike counts. Therefore, the heatmap showed a wide parameter space over which paired stimuli elicited combination responses, demonstrating that the CSN integrates inputs robustly across diverse temporal configurations. Together, these results extend our earlier findings by showing that while the first stimulus serves primarily as a temporal initiator of integration, the second stimulus strongly shapes the magnitude of CSN output. This division of functional roles highlights the robustness and flexibility of the network in processing syllable sequences.

### Stimulus strength

For stimulus intensity, we applied the same single-parameter variation strategy, systematically altering the intensity of one stimulus over a defined range while keeping the other fixed at its default value (Figure 7B). For the first stimulus (pale lavender trace), a minimum intensity of ∼100 pA is required to activate the network. Increasing this value to ∼140 pA resulted in a slight decrease in spike count, after which responses remained largely stable across the tested range. By contrast, the second stimulus (yellow trace) exhibited a similar activation threshold of 100 pA but exerted a much stronger influence on CSN activity. As S2 intensity increased, spike output increased, reaching a maximum at ∼400 pA. These results support the interpretation that the excitatory second stimulus plays a decisive role in shaping CSN responses, primarily through postsynaptic facilitation.

The two-dimensional intensity analysis (Figure 7D) further reinforced this pattern. Both stimuli exhibited a minimum intensity threshold for effective combination, as indicated by the absence of CSN spiking in the darkest regions of the plot. Increasing S1 intensity (y-axis) yielded little to no change in spike count, whereas increasing S2 intensity (x-axis) led to a clear progression from darker to lighter shades, reflecting higher spike counts. Together, these findings highlight the asymmetric contributions of the two inputs: while the first stimulus reliably initiates temporal integration, the second stimulus provides the critical excitatory drive that determines the magnitude of CSN output. In doing so, they provide a functional parameter map for input intensity combinations that support robust integration at the CSN level, highlighting the network’s capacity to generalize across a wide range of input strengths.

## DISCUSSION

Neural circuits across species and sensory systems exhibit a fundamental property, a temporal selectivity for specific combinations of different stimuli (Glass and Wollberg 1983, Margoliash 1983, Suga, O’Neill et al. 1983, McKenna, Weinberger et al. 1989). In this study, we developed a biophysically realistic network model to investigate the mechanisms underlying combination sensitivity. By embedding a temporal delay loop within a simplified circuit architecture, we demonstrated how sequential auditory inputs are dynamically transformed into an all-or-none facilitatory response at the CSN. Our results provide a mechanistic explanation for how spectrally distinct syllables can be temporally integrated over extended durations and then funneled into a narrow coincidence detection window, thereby enabling selective recognition of syllable sequences.

A key finding is that successful combination sensitivity relies on the obligatory transition between two distinct yet interdependent temporal coding windows: a prolonged integration window, spanning hundreds of milliseconds, and a brief coincidence detection window, spanning tens of milliseconds. The integration window is mediated by reverberatory activity in the delay loop and preserves a “memory trace” of the first stimulus. The coincidence window, in turn, is generated by intrinsic rebound dynamics of the CSN, which allow the excitatory input from the second stimulus to drive spiking only within a restricted temporal window. This sequential mechanism accounts for the experimentally observed all-or-none nature of CSNs (Margoliash 1983, Lewicki and Konishi 1995, Mooney, Rosen et al. 2002, Prather and Mooney 2008) and explains why reversing the order of stimuli abolishes facilitation.

Our network model is built upon concepts from early theories of synaptic superposition (Margoliash 1983, Lewicki and Konishi 1995) and postsynaptic priming (Mooney 2000, Mooney, Rosen et al. 2002, Prather and Mooney 2008) reported in the songbird literature, as well as support the longstanding hypothesis that postsynaptic facilitation underlies combination sensitivity in songbirds (Fitzpatrick, Kanwal et al. 1993, Casseday, Ehrlich et al. 1994, Sayegh, Aubie et al. 2011, Wenstrup, Nataraj et al. 2012). Specifically, the inhibitory input initiated by the first syllable does not directly drive CSN firing but instead hyperpolarizes the membrane potential, deinactivating T-type calcium channels. This sets the stage for a rebound response that can amplify subsequent excitatory input. The facilitation observed when the second syllable arrives within this rebound window highlights the critical role of intrinsic ionic conductances in shaping network-level computations. Perhaps the most critical role in our network design is played by the interneuron that is gating this network-level transition between the temporal integration modes. By dynamically modulating the level of inhibition, we showed that interneurons play a critical role in the response properties of CSNs. Our parameter analyses further show that while both T-type Ca^2+^ and H-currents contribute, the T-type conductance is indispensable for enabling rebound bursting. In mustached bats, postsynaptic facilitation is a prominent model explaining the spectrotemporal integration mechanism that govern the delay-tuned neurons’ sensitivity to pulse-echo frequency components (spanning sub-octave to multi-octave ranges), often occurring through coincidence detection of a delayed excitatory input with stimulus offset activity, mediated by rebound excitation following inhibition (Narins and Capranica 1980, Saitoh and Suga 1995, Aubie, Becker et al. 2009).

By systematically varying stimulus duration and intensity, we demonstrated that the model exhibits robust combination sensitivity across a wide range of syllable-like inputs. While the first stimulus primarily serves as a temporal initiator through loop activation, the second stimulus strongly determines the magnitude of CSN spiking. This asymmetry provides the network with both stability (insensitivity to variations in S1 duration or intensity) and flexibility (graded sensitivity to S2 features). Such properties may be particularly advantageous in natural song recognition, where variability in syllable duration, amplitude, and spectral content is common. Thus, the key to successful combination sensitivity lies not solely in the specific spectral properties of the chosen syllables but rather in the complex interplay of their temporal and spectral domains. These findings support our broader aim which is to develop a generalized framework for understanding temporal coding and combination sensitivity across diverse sensory systems.

The present model proposes a mechanistic framework in which temporal integration and postsynaptic facilitation cooperate to transform sequential auditory inputs into selective combination-sensitive responses. By incorporating both circuit-level reverberations and intrinsic cellular properties, our findings suggest that sequence sensitivity is not the product of a single synaptic motif, but rather an emergent property of the coordinated interaction between inhibitory and excitatory pathways. This resonates with prior in vivo reports of temporally precise syllable-specific facilitation in HVC (Margoliash and Fortune 1992, Doupe 1997, Suga 2015, Medina and Margoliash 2024). More broadly, our results highlight the importance of distributed network mechanisms for auditory temporal coding, bridging theories of coincidence detection with longer-timescale working memory processes.

## METHODS

Combination sensitivity is a multifaceted mechanism that is shared by various neurons and in different regions of the brain (Fuzessery and Feng 1983, Margoliash 1983, Metzner and Heiligenberg 1991, Olsen and Suga 1991, Sutter and Schreiner 1991, Margoliash and Fortune 1992, Tian and Rauschecker 1998, Carlson and Kawasaki 2004, Kanwal and Rauschecker 2007). Understanding the neural basis of this feature necessitates an investigation of network dynamics and the intrinsic cellular properties that could potentially shape neuronal firing patterns.

Computationally, to design our mathematical model in such a way to illustrate the effects of intrinsic and synaptic properties, we focused on the HVC of songbirds for three reasons: 1) a subset of HVC neurons are well-known to be combination sensitive (Margoliash and Fortune 1992, Theunissen and Doupe 1998, Mooney 2000), 2) the synaptic (Mooney and Prather 2005, Rosen and Mooney 2006) and intrinsic (Kubota and Taniguchi 1998, Daou, Ross et al. 2013) properties of neuronal population within the HVC nucleus are well described and 3) we can convey the underlying neurophysiological processes that governs combination sensitivity with an integrated set of excitatory/inhibitory synaptic currents and a set of inward/outward ionic currents. In other words, the excitatory – inhibitory interactions in our proposed network are modeled via biophysically realistic synaptic currents (AMPA/GABA) and the intrinsic properties that govern the neurons’ firing patterns are also modeled via biophysically realistic ionic currents that had been shown to be expressed pharmacologically in the HVC nucleus, making the underling discussion more realistic.

We developed single-compartment conductance-based Hodgkin-Huxley-type (HH) biophysical model of HVC neurons and connected them together via biologically realistic synaptic currents. Our model included excitatory HVCx-projecting neurons (HVCx) and local inhibitory interneurons (HVC_INT_). Since combination-sensitive neurons (CSNs) are considered to be a subset of HVCx neurons (Margoliash 1983, Mooney 2000), we modeled them as such. However, we further explored their unique properties by focusing on specific connectivity, synaptic inputs, and intrinsic membrane properties, as we will detail in the section on the desired network activity. The functional forms of activation/inactivation functions and time constants were based on previous published mathematical neural models (Hodgkin and Huxley 1990, Destexhe, Babloyantz et al. 1993, Terman, Rubin et al. 2002, Wang, Liu et al. 2003, Dunmyre, Del Negro et al. 2011, Daou, Ross et al. 2013). The parameters that were varied were the maximal conductance values of some ionic currents that vary among the various neuronal subtypes (Daou, Ross et al. 2013), as well as the synaptic conductances relevant to the network model. Every model neuron is represented by ordinary differential equations for the different state variables as illustrated below.

Simulations of the model neurons and network were conducted using the ode23 numerical integrator in MATLAB (MathWorks). The source code for these simulations will be made available online at our lab’s website and on ModelDB. In places where we were conducting brute-force searches of parameters, we used a High-Performance Computing (HPC) source at the American University of Beirut (AUB). Specifically, we employed the ‘octopus’ hardware cluster, operating under the Linux (CentOS 7) operating system.

Batch jobs were executed on the ‘arza’ partition, which comprises 18 computer nodes, each equipped with 16 cores and 64 GB of RAM, interconnected via an InfiniBand network.

### Ion channels of model HVC neurons

Our model included Hodgkin-Huxley type (Na^+^ and K^+^) currents for action potential generation (Hodgkin and Huxley 1952), a high-threshold L-type Ca^2+^ current (*I*_*CaL*_) for sustained firing (Kubota and Saito 1991, Long, Jin et al. 2010, Daou, Ross et al. 2013), a low-threshold T-type Ca^2+^ current (*I*_*CaT*_) for post-inhibitory rebound firing (Daou, Ross et al. 2013), a hyperpolarization-activated inward current (*I*_*h*_) to capture the sag potential (Kubota and Saito 1991, Dutar, Vu et al. 1998, Kubota and Taniguchi 1998, Daou, Ross et al. 2013), and a leak current (*I*_*L*_) for passive membrane properties. For interneurons, we integrated a large magnitude of the delayed rectifier K^+^ current conductance allowing these neurons to undershoot the resting membrane potential as seen experimentally (Kubota and Saito 1991, Dutar, Vu et al. 1998, Kubota and Taniguchi 1998, Daou, Ross et al. 2013). Additionally, a small-conductance Ca^2+^-activated K^+^ current (*I*_*SK*_) was added only to HVCx neurons (and the CSN) to model their observed spike frequency adaptation (Daou, Ross et al. 2013).

The membrane potential for each of the HVCx and HVC_INT_ neurons, including the CSN modeled as an HVCx neuron, obeys the following equations (Daou, Ross et al. 2013):

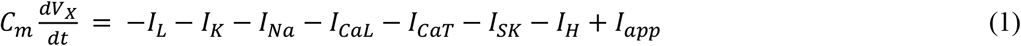

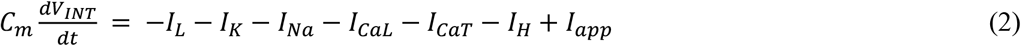

Where *C*_*m*_ is the membrane capacitance and *I*_*app*_ is the applied current. The associated equations and parameters for each of the activation/inactivation gating variables for each ionic current are given in (Daou, Ross et al. 2013) and shown below. Every single model of HVCx, and HVC_INT_ neuron had a total of 7 ODEs that govern their intrinsic dynamics. The integration of each synaptic current into a model neuron introduces an additional ODE to the system governing its membrane potential, as will be shown next. The values of the fixed parameter values are given in Tables S1-2.

### Voltage-gated ionic currents

The constant-conductance leak current is *I*_*L*_=*g*_*L*_(*V*−*V*_*L*_)

The remaining voltage-gated ionic currents have non-constant dependent currents with activation/inactivation kinetics:

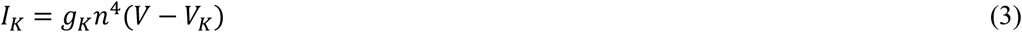

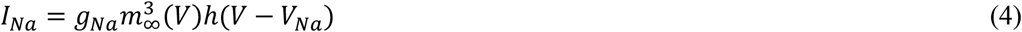

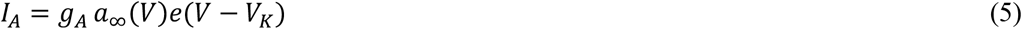

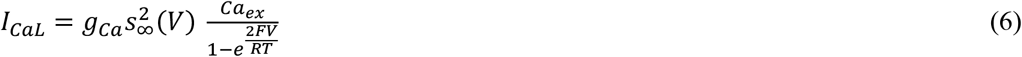

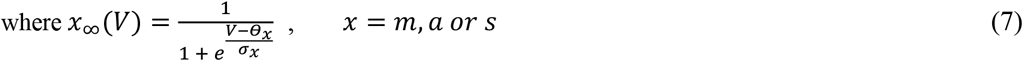

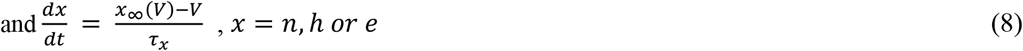

where *x*_∞_(*V*) for n and e is given by (7) and for *h*_∞_ as follows

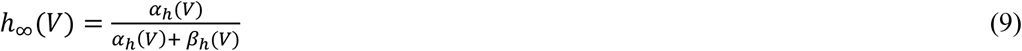

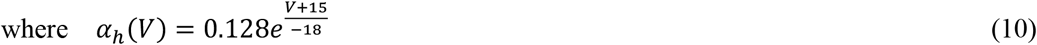

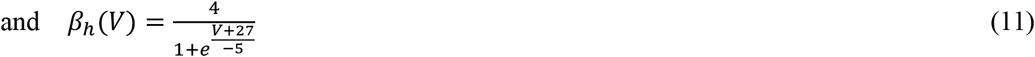

### Low-voltage activated T-type calcium current (I_CaT_)

The low-voltage activated T-type Ca^2+^ current is described by the Goldman-Hodgkin-Katz formula, with an expression similar to that given in (Terman, Rubin et al. 2002).

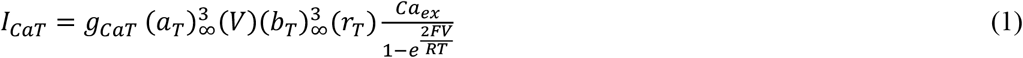

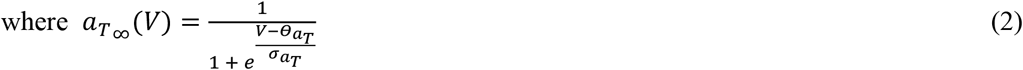

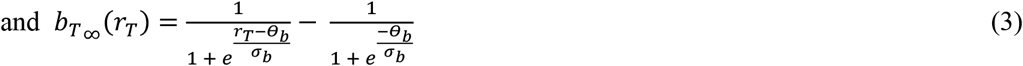

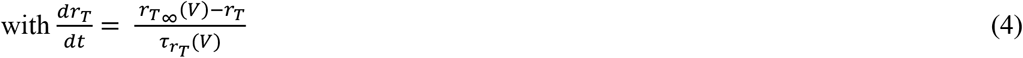

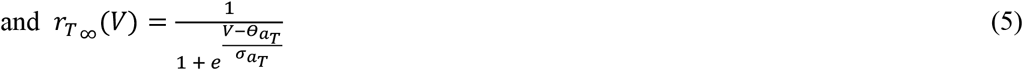

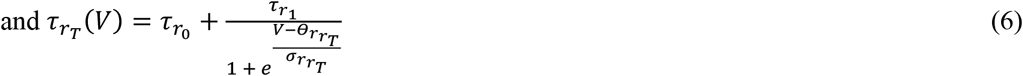

### Calcium-dependent potassium current (I_SK_)

The small conductance potassium current (*I*_*SK*_) is modeled as

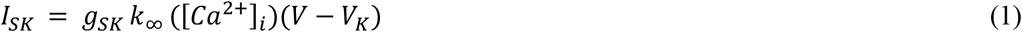

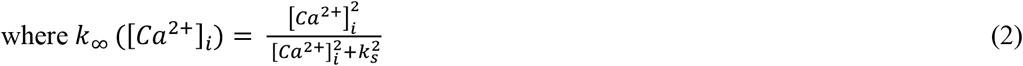

is the steady-state activation function of the SK current that is based on the levels of intracellular calcium. The constant *k*_*s*_ is the dissociation constant of the Ca^2+^-dependent current, and [*Ca*^2+^]_*i*_ is the intracellular concentration of free Ca^2+^ ions and is governed by:

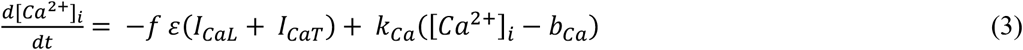

The constant *f* represents the fraction of free-to-total cytosolic Ca^2+^, whereas the constant ε combines the effects of buffers, cell volume, and the molar charge of calcium. Also, the constant *k*_*Ca*_ is the calcium pump rate constant, and *b*_*Ca*_ represents the basal level of Ca^2+^.

### Hyperpolarization-activated inward current (I_h_)

The hyperpolarization activated inward current’s activation is modeled as in (Destexhe, Babloyantz et al. 1993) using a fast component (*r*_*f*_) and a slow component (*r*_*s*_) as follows:

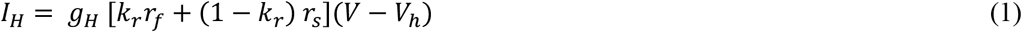

The fast activation component is given by:

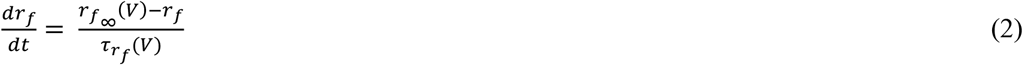

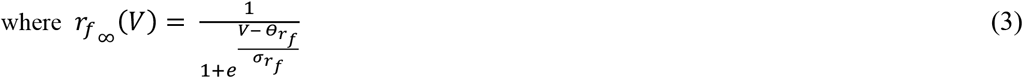

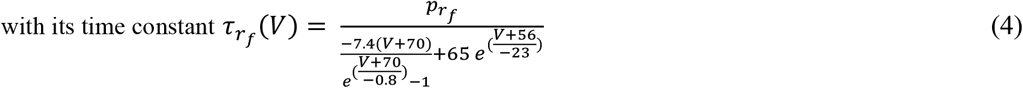

The slow activation component obeys:

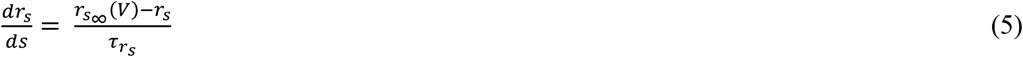

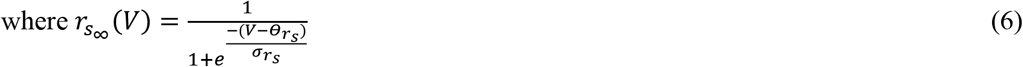

### Synaptic currents

To replicate the *in vivo* electrophysiological characteristics observed in HVC neurons, we integrated biologically realistic synaptic currents into the model alongside the above-described ionic currents. These synaptic currents represent excitatory (AMPA) and inhibitory (GABA_A_) synaptic inputs that are received by each presynaptic HVC neuron within the network. While there is a lack in experimental data onto the synaptic connections within the HVC in the case of combination sensitivity, we relied on data obtained from pharmacological dual synaptic connections from the broad network within the HVC, as described by (Mooney and Prather 2005, Kosche, Vallentin et al. 2015).

Each synaptic current represents the synaptic input(s) from the presynaptic cell(s) to the specific postsynaptic neuron, and is modeled as *I*_*syn*_ =∑_*X*_ *I*_*X*→*Y*_ where *I*_*X*→*Y*_ =*g*_*X*→*Y*_*s*_*X*→*Y*_(*V* − *V*_*X*→*Y*_). Here the summation is taken over the presynaptic HVC neurons where X represents a presynaptic cell, Y represents a postsynaptic cell, *V*_*X*→*Y*_ is the reversal potential for the synapse in the postsynaptic cell with *V*_*X*→*Y*_ =*V*_*AMPA*_ for excitatory input and *V*_*X*→*Y*_ =*V*_*GABA*−*A*_ for inhibitory input.

The model equations for the synaptic currents are the following, taken after (Destexhe, Mainen et al. 1994, Varela, Sen et al. 1997):

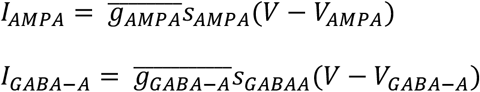

where *s*_*AMPA*_ and *s*_*GABAA*_ are given by

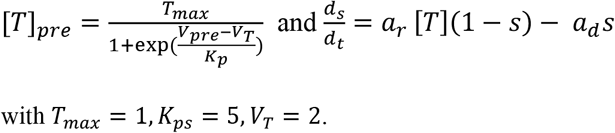

For GABA_A_, *a*_*r*_ =5 and *a*_*d*_ =0.18, while for AMPA, *a*_*r*_ =1.1 and *a*_*d*_ =0.19.

In our model, we incorporated only two synaptic currents: excitatory synapses (AMPA) and inhibitory synapses (GABA_A_). These AMPA receptor-mediated excitatory currents and GABA_A_ receptor-mediated inhibitory currents exhibit voltage-dependent behavior with relatively less complicated activation kinetics (Daou, Ross et al. 2013). Their well-defined dynamics are governed by established principles and do not require additional, intricate parameters for fine-tuning. We thus excluded NMDA and GABA_B_ receptor-mediated currents synaptic currents from our models for the following reasons: Both NMDA and GABA_B_ currents introduce additional layers of complexity due to their dependence on factors such as Mg^2+^ concentration (NMDA) and G-protein dynamics (GABA_B_) (Destexhe, Rudolph et al. 2001). Modeling these dependencies would necessitate incorporating supplementary ordinary differential equations (ODEs) and associated parameters that are currently lacking for HVC neurons. Our primary objective is to capture the core dynamics of excitation and inhibition within the network. AMPA and GABA_A_ currents can seamlessly represent these essential processes, enabling us to investigate the mechanisms underlying network behavior without introducing extra complexity.

### Stimulus presentation

While there is research reporting HVC’s afferent and efferent connectivity (Mooney 2000, Mooney and Prather 2005, Murphy, Lawley et al. 2020), little is known about the electrophysiological and anatomical identities of the auditory stimuli that the various classes of HVC neurons receive, their intrinsic and synaptic properties, and more importantly the mechanisms at which HVC neurons integrate this auditory information. In their model of song selectivity, (Drew and Abbott 2003) provided recorded songs as an input to their model in the form of spectrograms. This was passed through a set of linear filters, representing the action of neurons early in the auditory pathway, such that a spectrotemporal receptive field (STRF) function determined how the magnitude of the spectrogram at frequency f and a time period τ in the past, affects the neuron’s firing pattern. While it is a very interesting approach to model auditory stimulus as a spectrogram, the approach assumes a lot about how neurons process spectrotemporal information, putting an emphasis on the frequency feature in the spectral domain solely, and inherently masking critical information about the underlying stimulus complexity, probably limiting the visibility of other underlying neuronal mechanisms.

In our model, we assumed that when the first stimulus/syllable A is presented for T ms, an A-selective neuron is active during that T ms, and we modeled this selectivity by a simple DC-pulse that keeps the A-selective neuron firing during that T ms. We certainly considered the effects of stimulus intensity later on, but we believe this is not an unreasonable approach to modeling stimulus for the following reasons: 1) single syllable-selective neurons in the HVC are known to fire when the corresponding syllable is presented (Margoliash and Fortune 1992, Lewicki and Konishi 1995), i.e., when a syllable A or B is presented, we know that the underlying neuron is selective to these syllables primarily by their firing frequency (Lewicki and Konishi 1995, Lewicki 1996), 2) we are interested in the underlying intrinsic and interactive network mechanisms that occurs at different nodes within the network, which mainly occur after stimulus presentation. In other words, our main focus is on the network dynamics that shape and modulate the response firing of the CSN, irrespective of the specific representation of an “X” syllable. Whether the “X” syllable is represented by a neuron firing action potentials in response to a DC-pulse or a more complex current stimulus, the key understanding of the CSN activity relies primarily on different interactions between intermediate nodes within the network, rather than the input nodes themselves.

Therefore, every stimulus (syllable) presentation in our network is modeled as a DC-pulse applied to an HVC neuron. Unless otherwise specified, we set the stimulus duration to 50 ms, and the stimulus intensity to 200 pA, which generates a firing frequency of 200 Hz. We however explored extensively the effects of varying the stimulus duration and magnitude at various places in the results (Figure 7).

### Desired network activity

Our network model is built on the concept of postsynaptic priming (Mooney, Rosen et al. 2002, Rosen and Mooney 2003), a specific form of a well-established postsynaptic facilitation mechanism (Rosen and Mooney 2003, Ma, Jia et al. 2021). This mechanism provides an explanation for one form of spectrotemporal integration that is taking place at the level of the combination-sensitive neuron, thereby giving a certain mechanistic understanding behind one form of spectrotemporal integration at this level. This involves an inhibitory input followed by an excitatory input, where the precise timing of these inputs is crucial for triggering the desired integration. We have extended this mechanism by incorporating a circuit of upstream neurons that perform temporal integration prior to input convergence onto the combination-sensitive neuron. This circuit includes neurons that exhibit sustained activity during the temporal delay period between the arrivals of the two different sound elements (stimuli/syllables). The activity is generated through a feedback loop that is turned on by the onset of the first stimulus and then turned off at the onset of the second. Thus, we suggest that this temporally restricted activity within the upstream network effectively encodes temporal information during a specific integration window, reflecting the interval between the two stimuli, as found experimentally (Margoliash and Fortune 1992).

In general, combination sensitivity in higher-order auditory neurons is a highly selective neural process, governed by strict spectral and temporal conditions identified through experimental studies. Successful combination sensitivity is an all-or-none phenomenon, requiring the fulfillment of specific conditions. That is, in the case of a syllable pair AB, the CSN only responds when A precedes B. Reversed temporal order (BA) or single-syllable presentation (A or B alone) do not elicit a response. Thus, we focused on the intrinsic and synaptic parameters and their underlying mechanisms that play key roles in meeting these conditions and in governing the selective initiation of the process, maintaining the temporal propagation of information, triggering an active association between spectrally different stimuli, and performing the combination. While our primary focus was on the scenario where the first stimulus (syllable A) precedes the second stimulus (syllable B) within a specific temporal window, we also explored other scenarios where the desired behavior was not attained.

The assessment of the network’s ability to generate the desired firing behavior was done at two levels: 1) the CSN level, where the active association of stimuli occurs, representing spectrotemporal integration; and 2) the upstream network level, where initial and intermediate response mechanisms driven by presynaptic neurons influence and shape CSN responses. At the CSN level, the generated behavior was considered successful if it generated a burst of 4-12 spikes within a duration not exceeding 80 ms window. This burst should be triggered by the near-simultaneous convergence of selectively tuned inputs within a brief temporal window, termed the “coincidence window,” typically on the order of few milliseconds. Critically, this response is conditional upon both the presence and precise temporal order of the two stimuli. The CSN should remain subthreshold at all times when presented with either a reversed temporal order or by only one of the stimuli. At the presynaptic or upstream network level, specifically within the temporal delay circuit, desired behavior was characterized by the circuit’s ability to maintain patterned, continuous firing for the duration of the delay. This period of sustained activity, termed the “integration window,” can extend to several hundred milliseconds, consistent with a duration that revolves around 235 ± 73 ms for a syllable pair, as identified by (Margoliash and Fortune 1992). Critically, at the end of this temporal integration window, the collective output of the delay circuit should facilitate the active association of stimuli at the CSN level. Therefore, in addition to the CSN-level criteria described above, the network’s behavior was considered “desired” if it also passed this temporal-delay test. In our model, we ensured that the CSN exhibits robust firing when the delay between the two stimuli varies within this physiologically relevant range. Across all simulations, we carefully monitored the firing behaviors of all neurons in the network, ensuring that each neuronal population exhibited its specific intrinsic properties such as spike frequency adaptation, sag potentials, rebound firing, and others (Daou, Ross et al. 2013).

### Intrinsic and Synaptic conductance variations

Automated variations of model neuron conductances were performed in MATLAB to explore the parameter ranges for synaptic and intrinsic properties that maintain the network’s firing behaviors and mechanisms. At this step, we aimed to assess the network’s robustness and identify key parameters that enable the precise interactions and temporal alignment of neural events, as discussed in the “desired network activity” section, which are crucial for generating the desired CSN-level behavior; that is a successful combination of stimuli.

We employed a one-at-a-time (OAT) sensitivity analysis approach (Dey and Dimitrov 2023), wherein the effect of varying a single conductance parameter on network behavior was systematically assessed while all other conductances were held constant. We began by establishing a set of default intrinsic and synaptic conductance values for each neuron in the network, calibrated to generate the target behavioral output. Subsequently, for each conductance parameter, we conducted multiple simulation trials, introducing random perturbations around its baseline value. This iterative process was continued until deviations in network behavior were observed, allowing us to delineate the minimal and maximal bounds within which each conductance could vary without compromising the desired functional output.

During this exploratory phase, we excluded parameter regimes that produced biophysically unrealistic intrinsic properties in individual HVC neurons. Specifically, we discarded cases where HVC_X_ and HVC_INT_ neurons failed to exhibit characteristic features such as membrane sag, post-inhibitory rebound spiking, and depolarization-induced bursting. We also excluded simulations where spike amplitude and morphology were inconsistent with empirical observations. These included cases where HVC_X_ neurons failed to generate plateau-riding spikes or where HVC_INT_ neurons were lacking appropriate after-hyperpolarization following action potentials. While we acknowledge that constraining the range of permissible conductance values inherently limits the parameter space explored, this approach was necessary to ensure that all network behaviors remained within the bounds of known biophysical realism.

Given the critical role of inhibition and post-inhibitory rebound mechanisms in enabling successful stimulus combination in our network model, we concentrated our exploratory analysis on three intrinsic conductances: the small-conductance calcium-activated potassium current (*g*_*SK*_), the hyperpolarization-activated cation current (*g*_*h*_), and the low-threshold T-type calcium current (*g*_*CaT*_). All other intrinsic conductances were held constant throughout the simulations. Specifically, *g*_*Na*_ and *g*_*K*_ were fixed for each HVC neuron class based on values previously established to accurately reproduce spike morphologies, capturing features such as action potential upstrokes, downstrokes, and plateau potentials, under *in vitro* current-clamp conditions (Daou, Ross et al. 2013). We also fixed *g*_*CaL*_(L-type calcium conductance), as adjustments to *g*_*SK*_ alone are sufficient to maintain accurate fits to the observed spiking behavior.

Our iterative perturbation approach revealed that the fundamental combinatorial function of the network is relatively robust to variations in individual conductance values, indicating that precise tuning of parameters is not strictly required to sustain core network dynamics. The ranges of intrinsic conductance values that preserved network function are summarized in Table S3, and their distributions across simulations are visualized using boxplots in Figure S1A-C.

For clarity in network schematics and simulations, we adopted the following naming convention: inhibitory interneurons within *HVC*_*INT*_are designated with the letter “I” with subscripts used to differentiate individual neurons (e.g., *I*_1_ for interneuron 1). Excitatory projection neurons from the *HVC*_*x*_ class are labeled as “E,” again with corresponding subscripts. Additionally, neurons with specialized functional tuning are labeled “A,” “B,” and “CSN,” where “A” denotes neurons selectively responsive to the first syllable (stimulus 1 or S1), “B” to the second syllable (stimulus 2 or S2), and “CSN” identifies the combination-sensitive neuron.

Figure S1-A represents the distribution of T-type calcium conductance values for each of the following neurons in the network: E_1_, I_1_, I_2_, E_2_, and the CSN. Notably, these values exhibit a relatively narrow range of variation, between 0.1 to 4 nS, suggesting that these neurons are sensitive to specific levels of T-type calcium current. This sensitivity is crucial for maintaining their firing properties and overall network behavior. The CSN, in particular, required a more restricted range (0.4-0.6 nS) to ensure that it remains silent when only one stimulus is present, while still being able to respond to the appropriate temporal pattern of inputs.

The SK conductance values exhibited a slightly broader range of variability compared to the T-type calcium conductance (Figure S1-B). The CSN, in particular, had a more sensitive range of SK conductance values (0.1-3 nS). These values were chosen to ensure that the CSN firing remained within a biologically plausible range (maximum of 12 spikes per burst) and that other neurons in the network did not exceed 18 spikes per spike train.

The h-current conductance values for the CSN exhibited the widest range of variability, from 0.1 to 30 nS (Figure S1-C). This broad range reflects the importance of h-current in shaping the dynamic behavior of the CSN. We have shown in our results section that while T-type calcium currents are crucial for initiating the burst, h-current plays a key role in sustaining and prolonging the burst.

Although both currents contribute to depolarization after hyperpolarization, T-type calcium currents are more sensitive to specific temporal patterns of input.

A similar one-at-a-time (OAT) sensitivity analysis was performed for the synaptic conductance parameters, with the distinction that all synaptic conductances in the network were included in this evaluation. Each synaptic parameter was systematically varied around its default value while the others were held constant, following the same iterative procedure described for intrinsic conductances. This allowed us to determine the range of synaptic conductance values compatible with successful network function without compromising the fidelity of stimulus combination. The identified accepted ranges for synaptic conductances are summarized in Table S4, and their distributions across simulations are illustrated through boxplots in Figure S1D-F.

Each panel of the intrinsic boxplots focused on a specific group of synapses: those activated by the arrival (onset) of the first stimulus (Figure S1-D), those activated by the arrival of the second stimulus (Figure S1-E), and those sending their projections to the CSN (Figure S1-F). While the first two panels showed a wider range of variability in synaptic conductance values, the excitatory conductance from neuron B to the CSN had a more restricted range (1.4-3.6 nS). This limited range is necessary to ensure that the CSN does not fire when only one stimulus is present, as observed experimentally. This is illustrated in more detail in the Results section.

## SUPPLEMENTARY FIGURES

**Figure S1.**
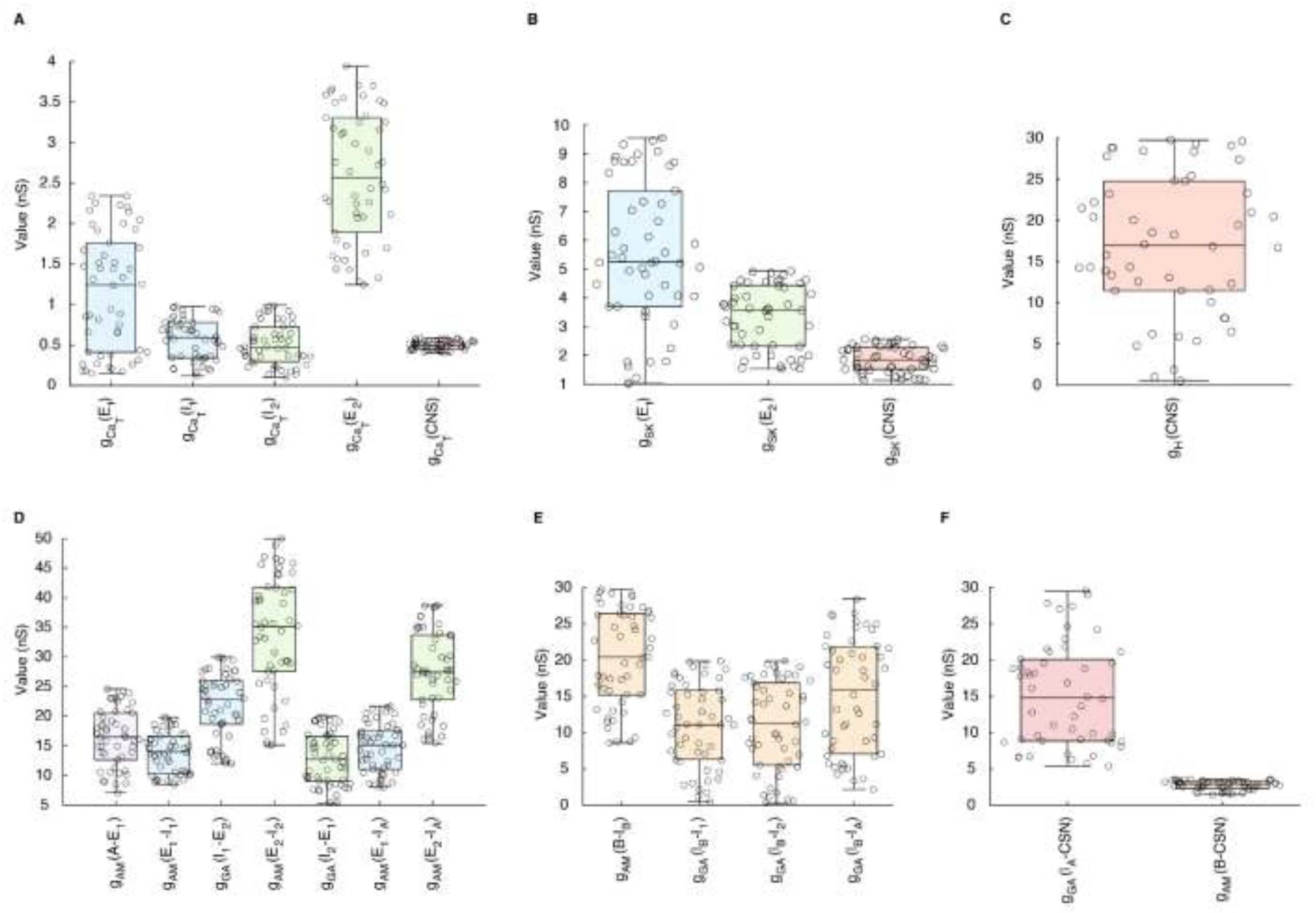
Distribution of identified conductance variability ranges for intrinsic ionic and synaptic currents for selected neurons from the network. The x-axis represents the neuronal population with the name of the neuron between brackets, and the y-axis represents the identified values of the sensitivity range of ionic current conductances. **(A)** Distribution of T-type calcium channel conductance (g_CaT_) for E_1_, I_1_, I_2_, E_2_, and CSN. **(B)** Distribution of calcium-activated potassium channel conductance (g_SK_) for E_1_, E_2_, and CSN. **(C)** Distribution of hyperpolarization-activated cyclic nucleotide-modulated current conductance (g_h_) for the CSN. **(D)** Boxplots representing synaptic conductance variability ranges for synaptic currents that are activated in response to stimulus 1, stimulus 2 **(E)** or arriving at the CSN **(F)**. Boxplots are color-coded according to their corresponding presynaptic neuronal populations within the network.

**Figure S2.**
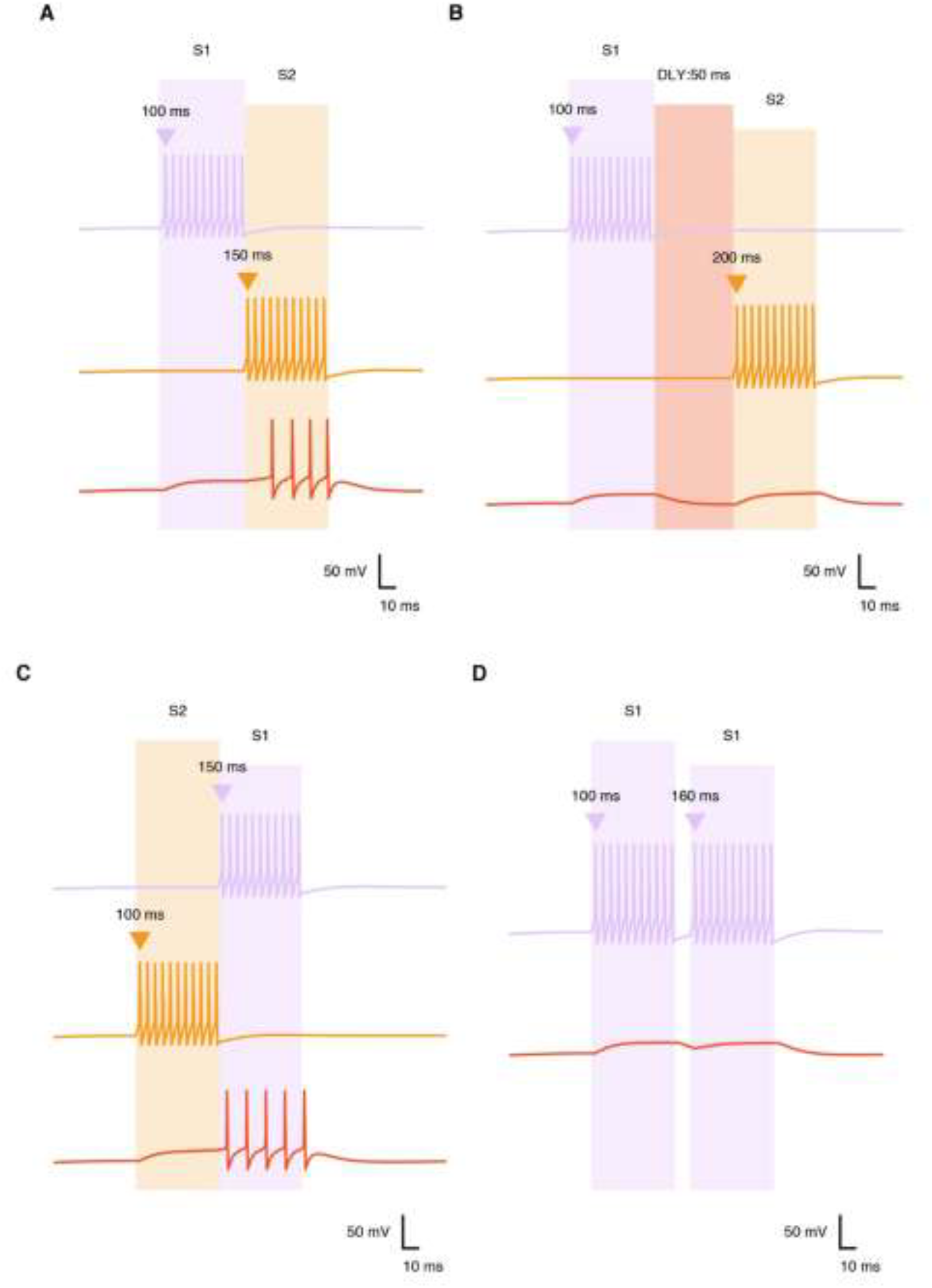
Simple network dynamics for the temporal summation scenario. Firing patterns for an excitatory neuron responding to the first stimulus (S1, pale lavender trace), an excitatory neuron responding to the second stimulus (S2, yellow trace) and the CSN (orange trace), shown under different conditions **(A-D)**. The onsets of S1 (pale lavender triangle) and S2 (yellow triangle) are varied across the different panels. Their respective durations are depicted by the shaded regions of the same colors. **(A)** S1 (50 ms) followed by S2 (50 ms)results in successful integration at the CSN level generating a subthreshold increase in the membrane potential in response to S1 which then surpass firing threshold at the onset of S2. **(B)** Temporal delay (DLY: 50 ms, soft red area) resulting in a dissociated subthreshold increase in membrane potential at the onset of both of S1 and S2, where the CSN goes back to resting state at offsets, thus resulting in a failure to associate at the CSN. **(C)** Reversed stimulus order generates a successful response, showing a spectral insensitivity. **(D)** Repeated stimulus presentation (in this case S1) did not succeed in elevating the membrane potential above firing threshold, resulting in subthreshold responses.

**Figure S3.**
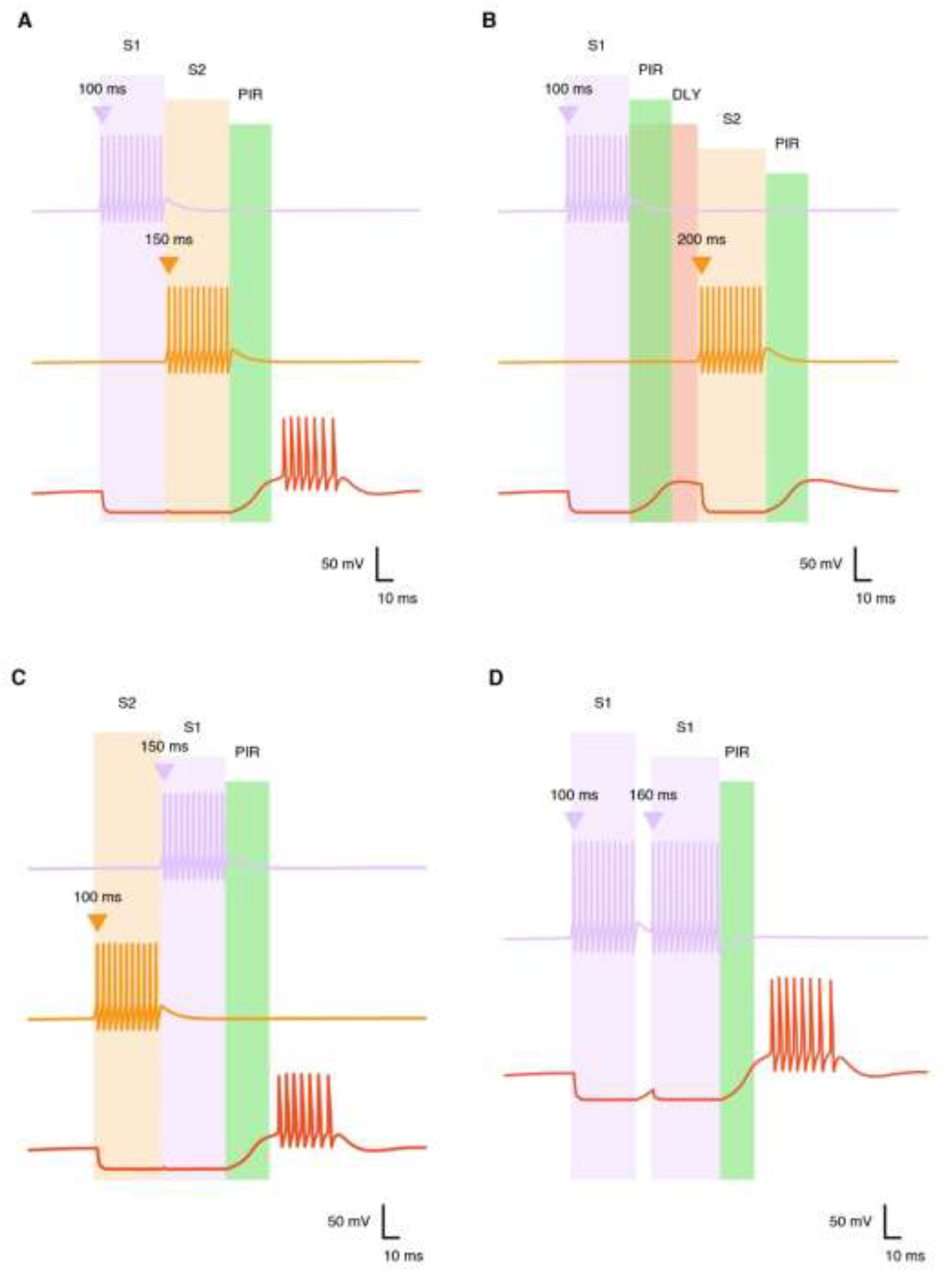
Simple network dynamics for the dual inhibitory scenario. Firing patterns for an inhibitory interneuron responding to the first stimulus (S1, pale lavender trace), an inhibitory interneuron responding to the second stimulus (S2, yellow trace) and the CSN (orange trace), shown under different conditions **(A-D)**. The onsets of S1 (pale lavender triangle) and S2 (yellow triangle) are varied across the different panels. Their respective durations depicted by the shaded regions of the same colors. **(A)** S1 (50 ms) followed immediately by S2 (50 ms) results in two successive membrane hyperpolarization of the CSN, the latter which initiates a post inhibitory rebound response (PIR, green shaded area) at the offset of S1-S2, leading to rebound depolarization and successful integration. **(B)** Temporal delay (DLY: shaded soft red area, 50 ms) between the offset of S1 and onset of S2 results in two dissociated PIR responses that did not reach firing threshold. **(C)** Successive reversed stimulus order and **(D)** repeated stimulus presentation (S1 in this case) result in successful response, similar to (A), showing spectral insensitivity.

**Figure S4.**
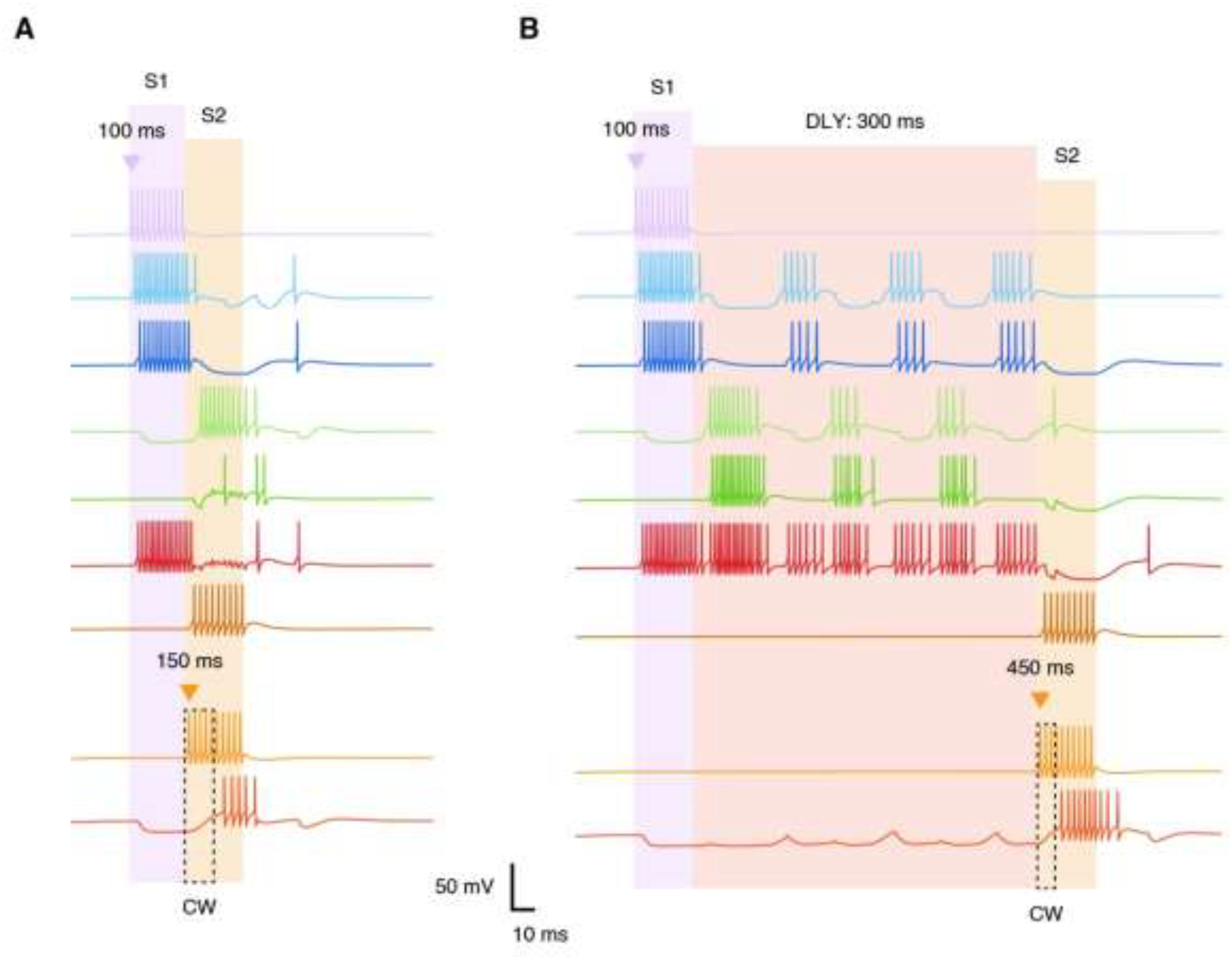
Combination-sensitive behavior at extreme delay durations of 0 ms and 300 ms. **(A)** Network dynamic in response to S1 followed by S2, showing neurons activated at the onset of S1 (pale lavender triangle), effectively hyperpolarizing the CSN’s membrane potential. The immediate onset of S2 (yellow triangle) at S1’s offset stops the temporal delay loop neurons from propagating their firing activity and opens a coincidence window (CW, dashed rectangle) at the level of the CSN, and successful integration. **(B)** Introducing an extended delay (DLY: 300 ms, shaded soft red area) between stimuli triggers same firing events at S1-S2 onset, leading to successful integration at the CSN level. Neurons of the temporal delay loop show a propagating and alternating activity.

**Figure S5.**
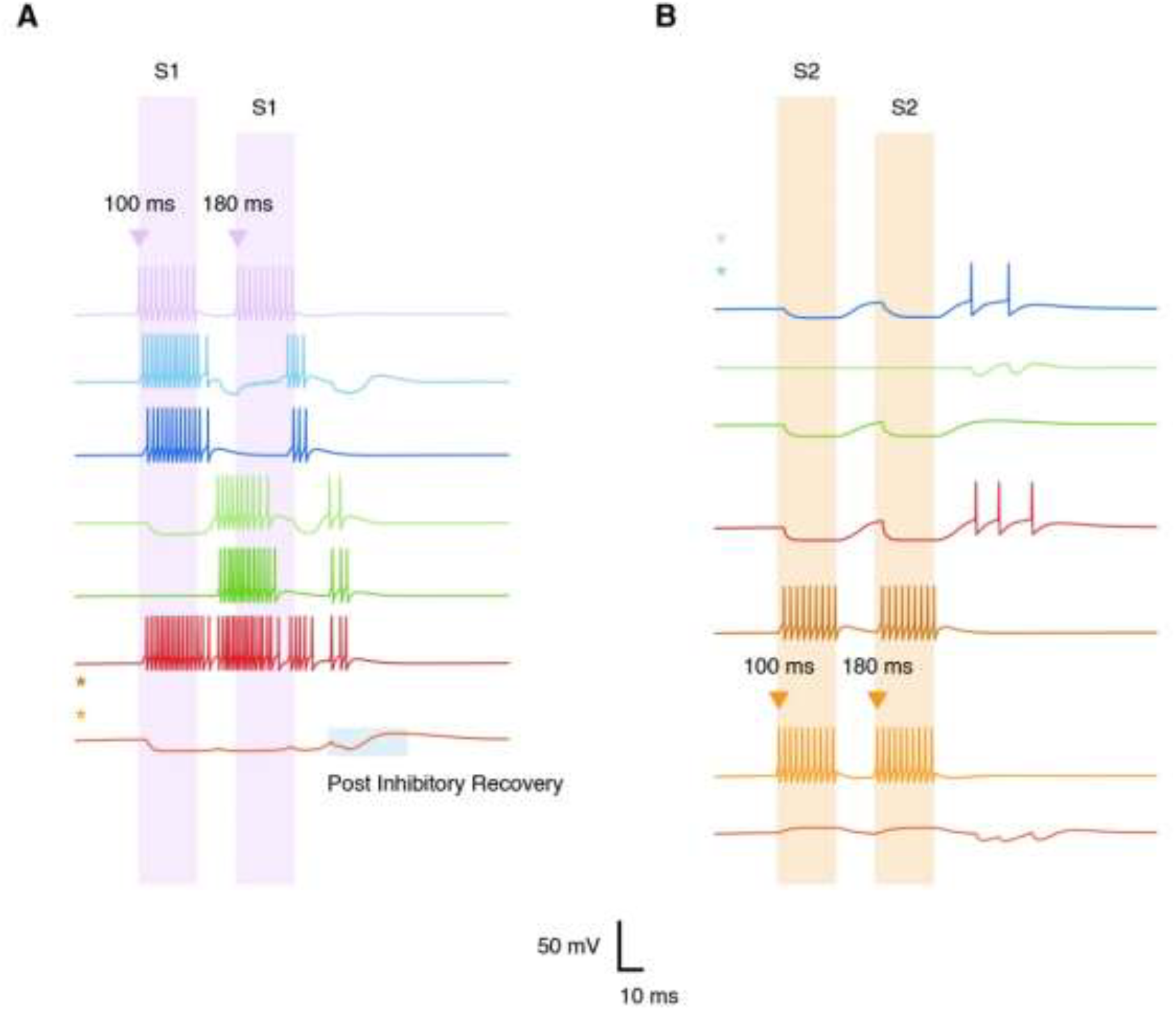
Spectral sensitivity of combination-sensitive neuron in response to repeated stimuli. **(A)** Repeated stimulus 1 presentation (S1) with a short delay duration (randomly chosen to be 80 ms, for visual presentation), results in triggering an inhibitory drive onto the CSN that stops gradually after the offset of the second S1, leading to a postinhibitory recovery, with no successful integration. Neurons of the temporal delay loop show a disrupted propagating activity. **(B)** Repeated stimulus 2 presentation (S2) with a short delay duration (randomly chosen to be 80 ms, for visual presentation), leading to two dissociated subthreshold depolarizing events at the CSN, which return to resting state after the offset of the second S2, with no successful integration. Untrigerred voltage traces are excluded, replaced with appropriate color-coded asterisks.

**Figure S6.**
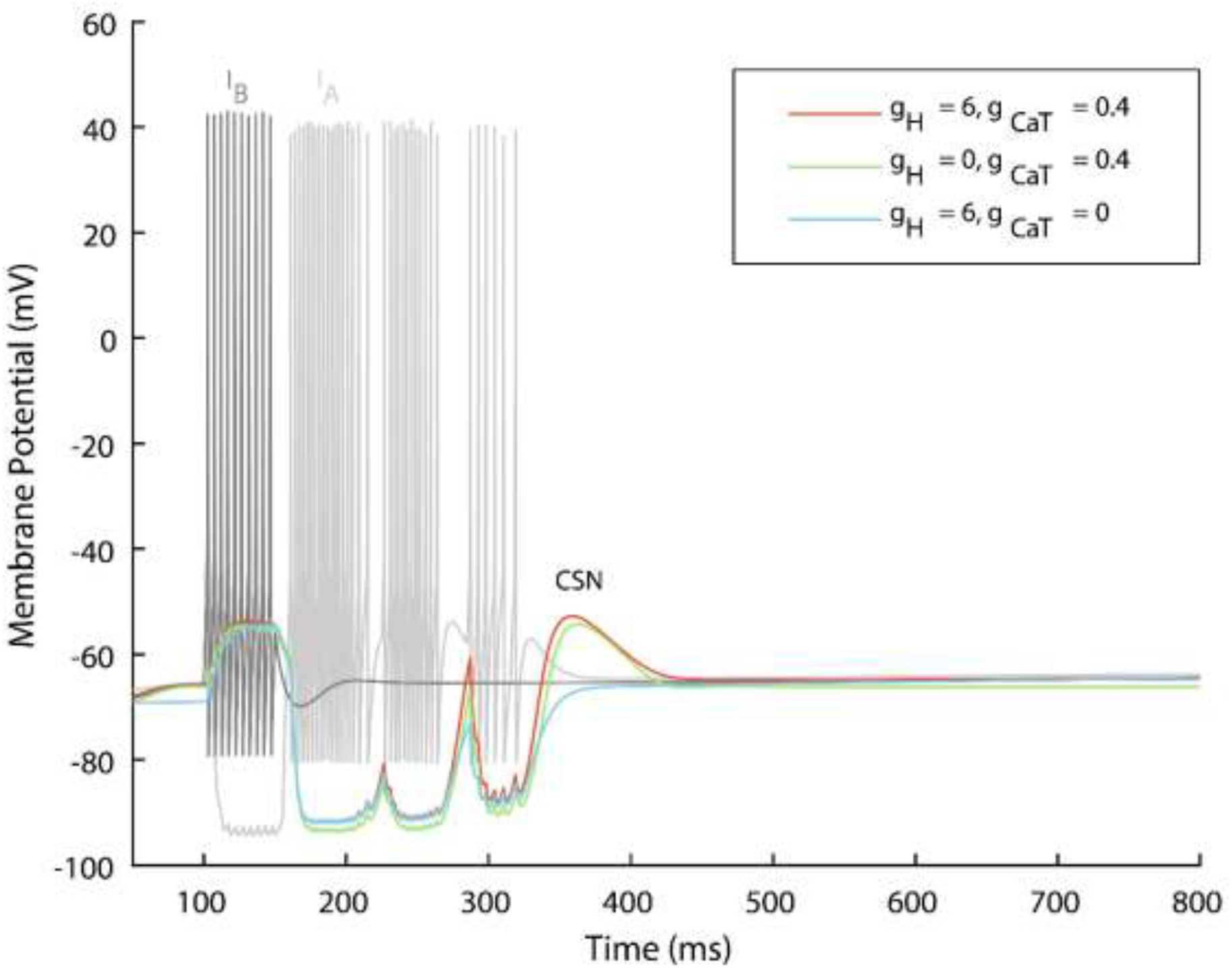
Temporal variation of stimuli and failed post-synaptic priming at the combination-sensitive neuron level. The figure illustrates voltage traces of a leading interneuron I_B_ (dark grey trace) representing stimulus 2 (S2), followed by interneuron I_A_ (light grey trace) representing stimulus 1 (S1) and their relevant effect on the membrane potential of the combination-sensitive neuron. Three traces of the combination-sensitive neuron (red, green, and blue) are shown to highlight the role of the intrinsic 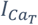 and I_H_ currents in response to conductance manipulation, with values shown in the legend. Red trace represents default conductance values. The green trace refers to blocking I_h_while keeping 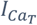 intact and the blue trace refers to blocking 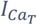 while keeping I_h_intact. In all cases, the association behavior is not attained.

## SUPPLEMENTARY TABLES

**Table S1.** Fixed parameter values used in all simulations.

| Parameter* | Value | Parameter | Value |
| --- | --- | --- | --- |
| $V_L$ | -70 mV | $V_K$ | -90 mV |
| $V_{Na}$ | 70 mV | $V_h$ | -30 mV |
| $g_L$ | 2 nS | $g_{Ca-L}$ | 19 nS |
| $g_{Na}$ | 800 nS | $\tau_{r_0}$ | 5 ms |
| $\tau_{r_1}$ | 15 ms | $\tau_n$ | 10 ms |
| $\tau_h$ | 1 ms | $\tau_{r_s}$ | 1500 ms |
| $\theta_m$ | -35 mV | $\theta_n$ | -30 mV |
| $\theta_s$ | -20 mV | $\theta_{r_f}$ | -105 mV |
| $\theta_{r_s}$ | -105 mV | $\theta_{r_{r_T}}$ | 68 mV |
| $\theta_{r_T}$ | -67 mV | $\theta_{a_T}$ | -85 mV |
| $\theta_b$ | 0.4 mV | $\sigma_m$ | -5 mV |
| $\sigma_n$ | -5 mV | $\sigma_s$ | -0.05 mV |
| $\sigma_{r_f}$ | 5 mV | $\sigma_{r_s}$ | 25 mV |
| $\sigma_{r_T}$ | 2 mV | $\sigma_{r_{r_T}}$ | 2 mV |
| $\sigma_{a_T}$ | -8 mV | $\sigma_b$ | -0.1 mV |
| $f$ | 0.1 | $\varepsilon$ | 0.0015 pA<sup>-1</sup> .μM. ms<sup>-1</sup> |
| $k_{Ca}$ | 0.3 ms<sup>-1</sup> | $b_{Ca}$ | 0.1 μM |
| $k_s$ | 0.5 μM | $p_{r_f}$ | 100 |

\* See text for parameter definitions.

**Table S2.** Parameter values that vary among neuron types.

| Parameter* | $HVC_x$ | $HVC_{INT}$ |
| --- | --- | --- |
| $g_K$ | 500 nS | 1200 nS |
| $g_h$ | 6 nS | 4 nS |
| $V_{RMP}$ | -75 mV | -67 mV |
| $C_m$ | 20 pF | 20 pF |
| $k_r$ | 0.3 | 0.01 |

\*See text for parameter definitions.

**Table S3.**
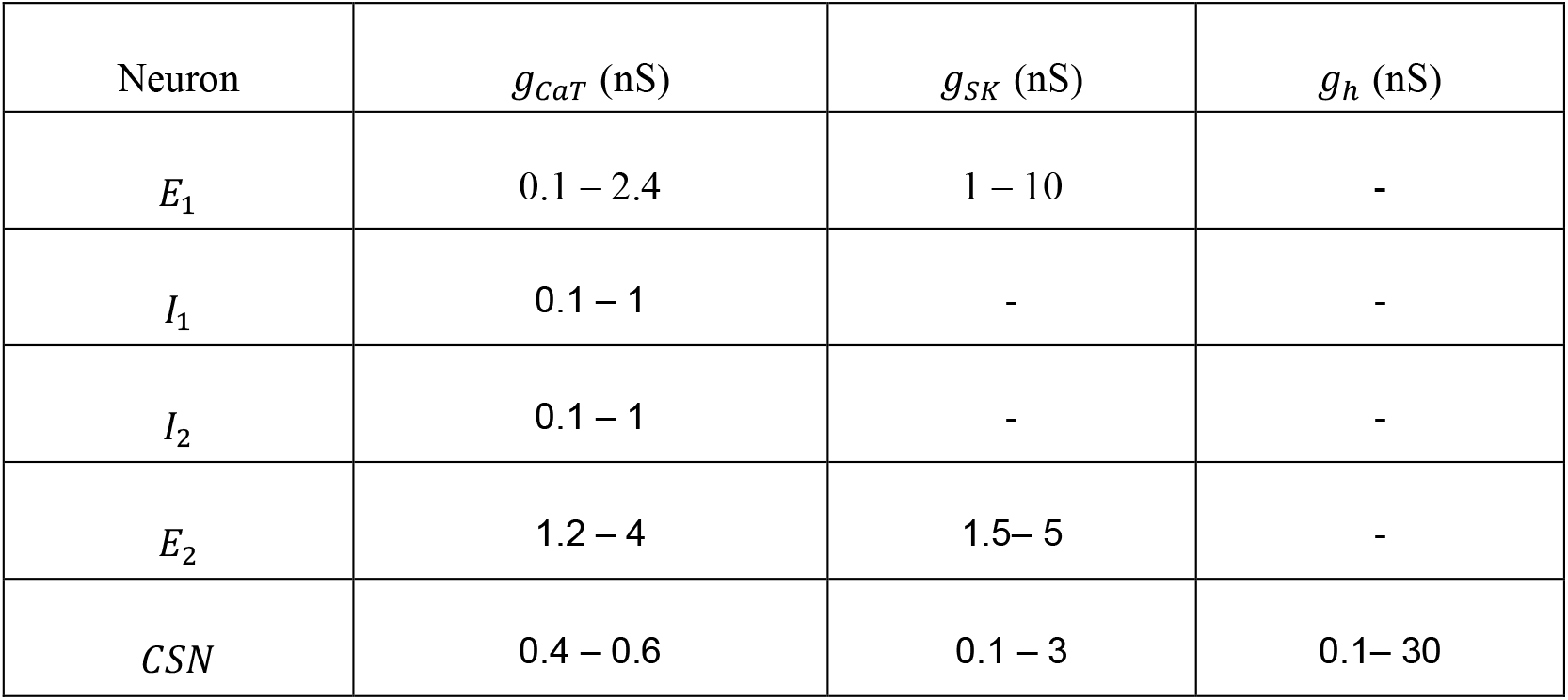
Identified ranges for key neuronal intrinsic conductances in the network.

**Table S4.** Identified ranges for synaptic conductances across the network.

| Synaptic Conductance values (nS) |  |  |  |
| --- | --- | --- | --- |
| $g_{AMPA}$ -Excitatory | | $g_{GABA}$ - Inhibitory | |
| $A \rightarrow E_1$ | 7 – 25 | $I_1 \rightarrow E_2$ | 12 – 30 |
| $E_1 \rightarrow I_A$ | 8 – 20 | $I_2 \rightarrow E_1$ | 5 – 20 |
| $E_2 \rightarrow I_2$ | 12 – 50 | $I_3 \rightarrow CSN$ | 5 – 30 |
| $E_1 \rightarrow I_3$ | 8 – 22 | $I_4 \rightarrow I_1$ | 0.1 – 20 |
| $E_2 \rightarrow I_3$ | 15 – 40 | $I_4 \rightarrow I_2$ | 0.1 – 20 |
| $B \rightarrow I_4$ | 8 – 30 | $I_4 \rightarrow I_3$ | 2.1 – 30 |
| $B \rightarrow CSN$ | 1.4 – 3.6 | | |

